# Diversification of Reprogramming Trajectories Revealed by Parallel Single-cell Transcriptome and Chromatin Accessibility Sequencing

**DOI:** 10.1101/829853

**Authors:** Qiao Rui Xing, Chadi El Farran, Pradeep Gautam, Yu Song Chuah, Tushar Warrier, Cheng-Xu Delon Toh, Nam-Young Kang, Shigeki Sugii, Young-Tae Chang, Jian Xu, James J. Collins, George Q. Daley, Hu Li, Li-Feng Zhang, Yuin-Han Loh

## Abstract

To unravel the mechanism of human cellular reprogramming process at single-cell resolution, we performed parallel scRNA-Seq and scATAC-Seq analysis. Our analysis reveals that the cells undergoing reprogramming proceed in an asynchronous trajectory and diversify into heterogeneous sub-populations. BDD2-C8 fluorescent probe staining and negative staining for CD13, CD44 and CD201 markers, could enrich for the *GDF3*+ early reprogrammed cells. Combinatory usage of the surface markers enables the fine segregation of the early-intermediate cells with diverse reprogramming propensities. scATAC-Seq analysis further uncovered the genomic partitions and transcription factors responsible for the regulatory phasing of reprogramming process. Binary choice between a FOSL1 or a TEAD4-centric regulatory network determines the outcome of a successful reprogramming. Altogether, our study illuminates the multitude of diverse routes transversed by individual reprogramming cells and presents an integrative roadmap for identifying the mechanistic part-list of the reprogramming machinery.

## INTRODUCTION

Somatic cells can be reverted to pluripotency by inducing the expression of four transcription factors namely OCT4, SOX2, KLF4 and MYC in a process known as cellular reprogramming^1–6^. Discovery of this phenomenon has raised the hopes for advancing the field of regenerative medicine^7^. However, cellular reprogramming suffers from extremely low efficiency especially for the human cells, resulting in a heterogeneous population in which few cells can be characterized as pluripotent^8–10^. Although a handful of studies analyzed bulk population to understand the reprogramming mechanisms^11–17^, ensemble measurement of the heterogeneous population impedes the discerning of transcriptomic and epigenetic changes taking place in the minority of cells undergoing the route towards successful reprogramming. Single-cell sequencing technologies provide tools to decipher the types of cells present in a heterogeneous mixture^18^. In the present study, we adopted the parallel genome-wide single-cell assays of scRNA-Seq and scATAC-Seq^19–21^ to profile transcriptome and chromatin accessibility of human reprogramming cells across various stages. We identified cellular diversification and trajectories where individual cell displays different dynamics and potential for reprogramming. Moreover, with a set of cell surface markers and a fluorescent probe BDD2-C8, we were able to enrich for the early-intermediary cells undergoing route towards successful reprogramming. In addition, we identified the modulators driving the changes in gene regulatory network and chromatin accessibility, as cells advanced towards the diverse reprogramming trajectories. Of note, the pivot from FOSL1 to TEAD4-centric regulatory networks is essential for the acquisition of the pluripotent state.

## RESULTS

### Single-cell Profiling of Cell-Fate Reprogramming

To study the heterogeneity of human reprogramming, we analyzed a total of 33468 scRNA-Seq and scATAC-Seq libraries with good quality, including day 0 (BJ), day 2 (D2), day 8 (D8), day12 (D12) and day 16 (D16) OSKM-induced reprogramming cells (Figure 1a and Supplementary Table 1). On D16, cells were sorted using the TRA-1-60 marker to distinguish the successfully reprogrammed (D16+) from the non-reprogrammed (D16-) cells (Figures 1a and S1a-b). The generated iPSCs were characterized with immunostaining, DNA methylation, and terotoma assay (Figures S1c-e). Two distinct approaches were used for scRNA-Seq library preparation. Microfluidic cell capture-based assay (Fluidigm C1) reads the full-length transcripts from hundreds of cells with high resolution, whereas the droplet-based assay (10X Genomics) probes the 3’ end of the transcripts from thousands of cells albeit at a relatively low genomic coverage. In addition, we have also screened for fluorescence probes to distinguish early-intermediate cells poised for successful reprogramming (Figure 1a). The cumulative data enable us to characterize the subpopulations in-depth and to construct a trajectory map of human reprogramming.

**Figure 1.**
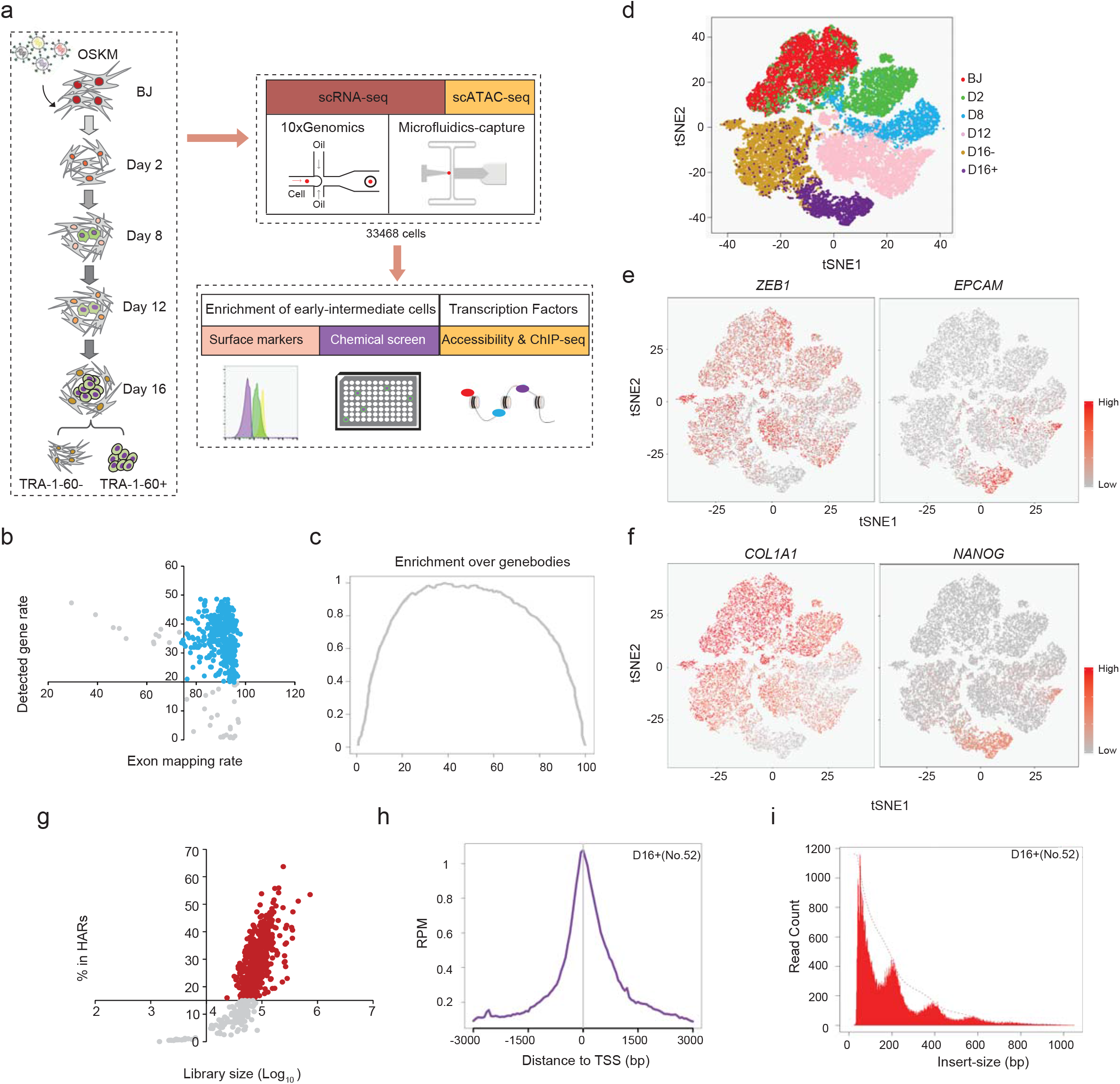
Schematic of the single-cell systems used for de-convoluting the heterogeneity in human cellular reprogramming. (a) Overview of the prepared single-cell NGS libraries at the indicated time-points of the human cellular reprogramming. Two single-cell platforms were utilized for the study. Microfluidics platform was used to prepare 439 single cell RNA-Seq and 891 single cell ATAC-Seq libraries (Duplicates). 10x Genomics platform provided us with an additional 32138 cells for scRNA-Seq analysis. (b) Quality control of Microfluidics-capture based scRNA-Seq libraries. Dotplot demonstrates the exon mapping percentage (X-axis) of each scRNA-Seq libraries, along with its corresponding detected gene rates (Y-axis). Axis crosses each other at the cutoff values that filter the libraries. Blue dots represent libraries that passed the QC filters. (c) Average enrichment of scRNA-Seq libraries over genebodies. (d) t-SNE plot of the prepared 10x scRNA-Seq libraries based on total cellular transcriptomes. (e-f) Super-imposition of the single-cell expression levels of MET genes: *ZEB1* and *EPCAM* (e), and fibroblast and pluripotent genes: *COL1A1* and *NANOG* (f), on the tSNE plot. The expression ranges from no expression (grey) to high expression (red). (g) Quality control of scATAC-Seq libraries. Dotplot demonstrating the library size (X-axis) of each scATAC-Seq library, along with its corresponding contribution to its respective time-point’s HARs (Y-axis). Red dots represent the cells that pass the QC filters. (h) Average enrichment profile of a D16+ scATAC-Seq library around Transcription Start Sites (TSS) of the genome with a window of −3K to 3K. Y-axis denotes the average normalized read counts of the library over the indicated region in the genome (x-axis). (i) Histograms of insert size metric of a D16+ scATAC-Seq library revealing a nucleosomal pattern, which is characteristic of a good ATAC-Seq library.

Majority of the capture-based scRNA-Seq libraries showed high exon mapping percentage (>=75%) and gene detection rate (>=20%) and displayed even distribution over the gene-bodies (Figure 1b-c, and Supplementary Table 1). Furthermore, epithelial and pluripotency genes were progressively expressed with the advancement of reprogramming, as opposed to the mesenchymal and fibroblast genes (Figure S1f). Similarly, majority of the 10X libraries were of good quality (Figures S1g-h). Notably, t-SNE plot revealed a dynamic transcriptomic transition from the parental BJ to D16+ cells (Figure 1d). Expectedly, *ZEB1* (mesenchymal) and *COL1A1* (somatic) were abundantly expressed in the early time-points and non-reprogrammed cells (Figures 1e-f). On the contrary, *EPCAM* (epithelial), and *NANOG* and *LIN28A* (pluripotent) were expressed highly in the successfully reprogrammed cells (Figures 1e-f and S1i). Likewise, most of the scATAC-Seq libraries passed the previously reported QC indices^19^, and exhibited enrichment over TSS regions and nucleosomal distributions (Figure 1g-i and S1j-k, and Supplementary Table 1). Collectively, we generated reliable libraries for tens of thousands of reprogramming cells, providing a rich resource to decipher its deep molecular mechanisms.

### Deciphering the Heterogeneous Subgroups with Diverse Reprogramming Potentials

Due to its high sensitivity, capture-based scRNA-Seq libraries were analyzed first to decipher the heterogeneity. CellNet^22^ revealed the dynamics of reduced fibroblast similarity and increased ESC correlation (Figure S2a). To determine the diverse populations present at each reprogramming time-point, we clustered scRNA-Seq libraries using Reference Component Analysis (RCA)^23^. Interestingly, BJ cells correlated significantly with the smooth muscle lineage, which was also detected across the published BJ libraries (Figures S2b-c). D2 cells were marked by four distinct subgroups (G1-G4), among which G1-3 cells displayed lower correlation to the fibroblasts and mesenchymal stem cells (MSCs) (Figures 2a and S2d). D8 cells were distributed among three discrete sub-populations (Figures 2b and S2d). D8 G1 cells corresponded to the fibroblasts, smooth muscles, myocytes, and MSCs lineages, whilst G3 cells displayed substantial similarity to the pluripotent stem cells (PSCs). Interestingly, D8 G2 cells represented the intermediate state. Two sub-populations were present in the D16+ cells (Figures 2c and S2d). D16+ G2 cells were highly associated with PSCs, while G1 cells maintained detectable correlation to the MSCs, adipose cells, and endothelial cells, other than PSCs. In summary, RCA analysis strongly indicates that the reprogramming cells are highly diverse, some of which may deviate from the route to pluripotency and acquire alternative lineage cell-fates.

**Figure 2.**
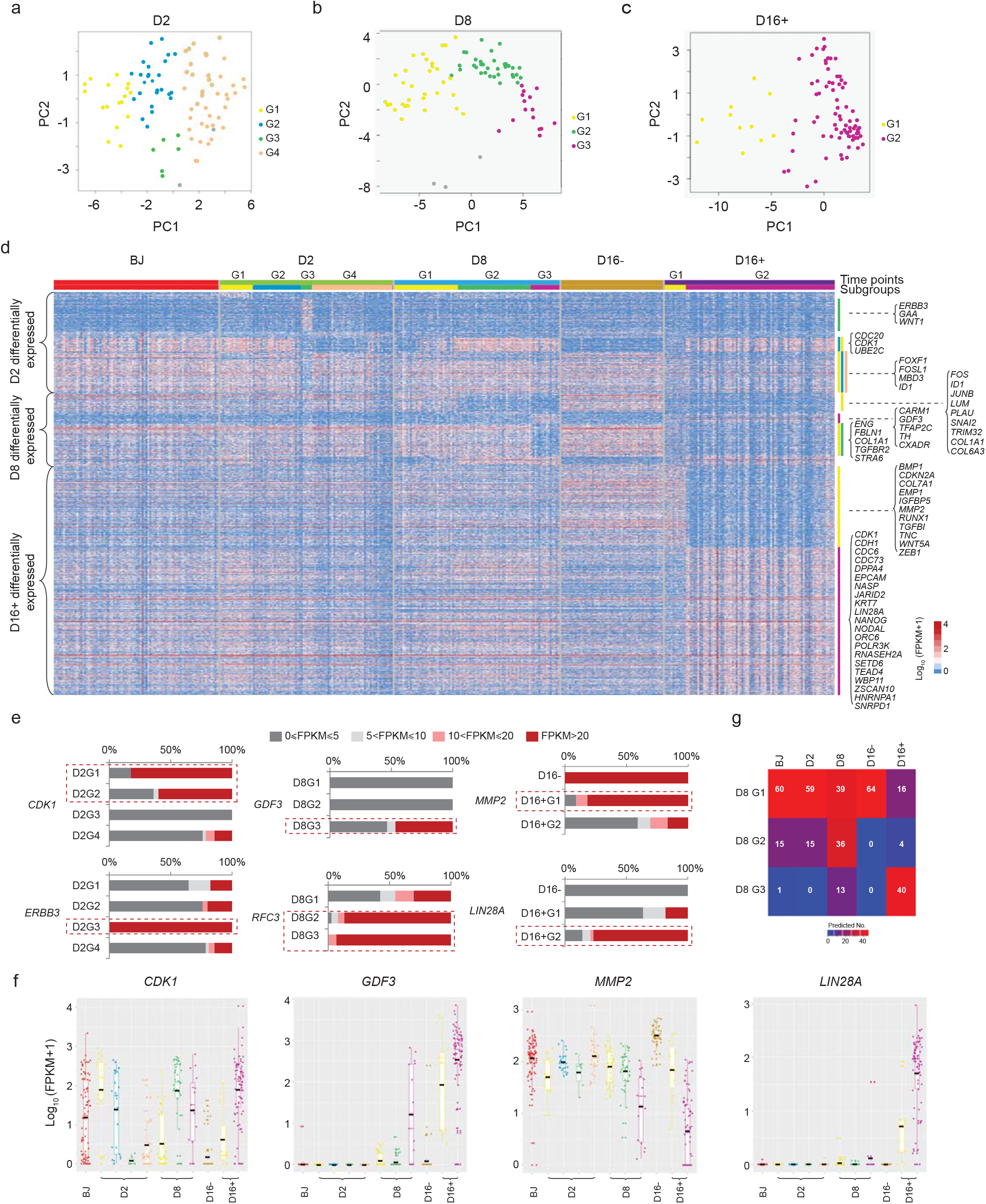
Identification of diverse reprogramming subgroups. (a-c) RCA clustering of D2 (a), D8 (b), and D16+ (c) cells based on the profile of the expressed genes. The PCAs show subgroups of reprogramming cells at individual time points. Each color represents a subgroup. (d) Heatmap showing dynamics of the genes differentially expressed among 4 subgroups of D2, 3 subgroups of D8, and 2 subgroups of D16+ cells, across the reprogramming process. Color represents the expression level, ranging from dark blue (low) to dark red (high). The expression values are Log_10_ (FPKM+1). The color code on top represents the time points (above) and their respective subgroups (below). (e) Bar charts demonstrating the expression pattern of D2 (left), D8 (middle), and D16+ (right) subgroup specific genes. Color represents the expression level. X-axis represents the percentage of cells with the expression within the indicated range. Red dotted boxes highlight the subgroups with high expression of the respective gene. (f) Box plots showing the single cell expression of the differentially expressed genes *CDK1* (D2), *GDF3* (D8), *MMP2* (D16+), and *LIN28A* (D16+) across all the time points and their respective subgroups. Lines in the box represent the median expression in the respective subgroups or time points. The expression values are Log10 (FPKM+1). (g) Confusion matrix of the scRNA-seq libraries of all time points, using Random Forest algorithm. Color represents the number of libraries from each time point predicted to be similar to the indicated subgroups of D8. Scale ranges from dark blue (low number) to dark red (high number).

We then performed differential gene expression (DGE) analysis to evaluate the subgroup specific genes (Figure 2d and Supplementary Table 2). Among D2 subgroups, G3 cells had the most distinct transcriptomic profile with exclusive expression of a remarkable number of genes, including *ERBB3* (Figures 2d-e). Majority of D2 G1-2 genes, for example *CDK1*, were expressed highly in D16+ G2, suggestive of their higher reprogramming propensity (Figures 2d-f). Among D8 subgroups, G1-2 specific genes were expressed highly in BJ and D16-but not the D16+ cells, such as *JUNB*, *LUM*, *COL1A1*, and *COL6A3* (Figure 2d). The opposite trend was observed for D8 G2-3 genes. *RFC3* was vastly expressed in D8 G2-3 cells, whereas *GDF3* was specifically expressed in the G3 (Figures 2d-f). In agreement with the correlation to the differentiated lineages, D16+ G1 specific genes, including *MMP2*, were also expressed highly in the D16-cells, suggesting that D16+ G1 cells were at most partially reprogrammed (Figures 2d-f). On the other hand, epithelial genes and pluripotent genes including *CDH1*, *NANOG* and *LIN28A*, as well as genes associated with mRNA splicing and transcription of the small RNAs including *WBP11* and *POLR3K*, were specifically expressed in D16+ G2 cells. Intriguingly, depletion of *POLR3K* and *WBP11* resulted in ablated reprogramming efficiency, indicating the functional importance of the D16+ G2 specific genes for reprogramming (Figure S2e). Additionally, D8 G3 and D16+ G2 specific genes exhibited high stemness score for PSCs, whereas D8 G1-2 and D16+ G1 specific genes significantly associated with the differentiated lineages (Figures S2f-g).

To test the notion of the diverse reprogramming potentials, D8 subgroups were correlated to the subgroups of various time-points (Figure S2h). Interestingly, G3 cells highly correlated with D16+ G2 cells, whereas G2 cells represented an intermediate state in which the cells were moderately correlated with all D16 cells. G1 cells, on the other hand, strongly correlated with D16- and cells from the early time-points (BJ and D2). In-house devised classifier displayed a similar correlation trend of the reprogramming time-points to the D8 subgroups (Figure 2g).

### Pseudotemporal trajectory of reprogramming cells

We next analyzed 10X libraries to construct pseudotemporal map of cellular reprogramming. CellNet and RCA of 10X libraries reproduced the reprogramming dynamics and diverse subgroups with variable lineage correlations (Figures S3a-c). Resultant pseudotemporal trajectories^24,25^ consisted of 9 states and 4 branching events (Figures 3a and S3d). Notably, pseudotime highly correlated with the reprogramming time-points (Figures 3a-c). We then traced the trajectory of RCA subgroups. Interestingly, majority of the D2 G1-3 cells were found in state 3 (95% of G1, 65% of G2, and 80% of G3), whereas G4 cells scattered across the early states (Figure S3e). Joint RCA analysis demonstrated that D2 G1-2 cells clustered closer to the D8 G2 cells and correlated stronger to the ESC fate, indicating their higher reprogramming propensity (Figures S3f-g). Of note, D8 G1 cells enriched across the different states other than state 9 (successfully reprogrammed) (Figures 3d-e). On the contrary, D8 G3 cells were mostly found in state 9. D8 G2 cells distributed across the intermediate (3-5) and late states (7-9). Intriguingly, state 4 comprised almost entirely of D8 G2 cells (544 vs. 627) (Figures 3d-e). Expectedly, D16+ G2 cells were the major constituents of state 9, whereas cells of D16+ G1 were enriched in both state 8 (non-reprogrammed) and state 9, corroborating their partially- and non-reprogrammed identities (Figures 3d and S3h). Further, subgroup specific markers were expressed differentially along the pseudotime axis (Figure 3f). *RFC3* (D8 G2-3) and *GDF3* (D8 G3), and *NANOG* and *LIN28A* (D16+ G2) were expressed highly in the cells on the successful reprogramming trajectory, whereas *MMP2* (D16+ G1) showed the opposite trend. Next, we determined the gene expression patterns and significant biological processes defining the trajectory (Figure S3i and Supplementary Table 3). At branching event 1, cells with high levels of lineage genes advanced towards successful reprogramming, whereas cells with abundant DNA replication genes deviated from the path (Figures 3g-h and S3i). Interestingly, successfully reprogrammed cells at branching point 4 were highly active for DNA replication and mRNA splicing, which were implicated to be essential for reprogramming^26,27^. On the other hand, collagen and extracellular matrix related genes emerged to be detrimental for reprogramming at the branching point 3 and 4, and lineage processes such as angiogenesis and epidermis development contributed to the unsuccessful reprogramming at branching point 4. Together, these analyses provide a plethora of data for identifying the novel processes and modulators affecting the reprogramming trajectory at an unprecedented single-cell resolution.

**Figure 3.**
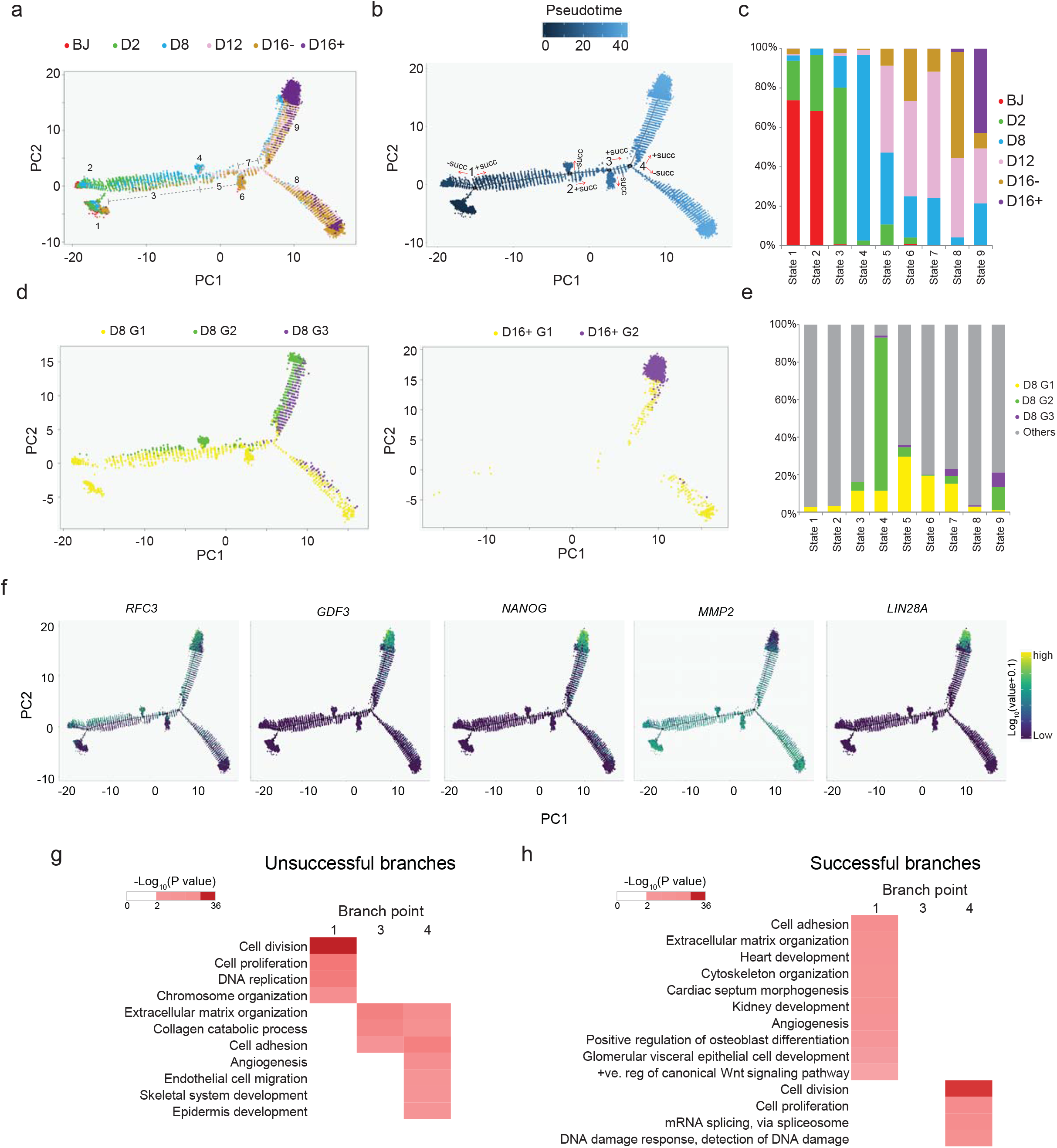
Trajectories of cellular reprogramming. (a) Trajectory of reprogramming cells identified from the 10x scRNA-Seq libraries based on DDRTree dimension reduction. Color represents the time points. (b) Pseudotime calculation based on the DDRTree trajectory. Color indicates pseudotime, ranging from dark blue (early) to light blue (late). The plot shows four branching events. “+succ” and “-succ” represents the successful and unsuccessful braches. (c) Stacked columns indicating the percentage of cells belonging to the respective time-points in the reprogramming trajectory states. Color represents the time points. (d) Super-imposition of D8 subgroups (left) and D16+ subgroups (right) on the trajectory of reprogramming. Color represents the subgroups. (e) Stacked columns revealing the percentage of cells of the respective D8 subgroups in the indicated reprogramming trajectory states. Colors represent the subgroups of D8 and grey color indicates cells of the other time points. (f) Superimposition of the expression of D8 subgroup genes (*RFC3*, *GDF3*) and D16+ subgroup genes (*NANOG*, *MMP2* and *LIN28A*) on the reprogramming trajectories. Expression ranges from purple (low) to yellow (high). (g-h) GO analysis of the differentially expressed genes associated with the unsuccessful (g) and successful (h) reprogramming branches. The enrichment score ranges from white (no) to red (high). The braches are labelled in Fig.3b.

### Toolkits to enrich for early-intermediate cells with diverse reprogramming potentials

To enrich for the intermediate cells with high reprogramming potential, we conducted a screen for a library of 34 DOLFA^28,29^ fluorescent dyes (Figure S4a). Candidate dyes differentially stain the intermediate reprogramming cells with the accelerated dynamics upon treatment with a TGFβ inhibitor^30,31^, A83-01. BDD1-A2, BDD2-A6, and BDD2-C8 were identified as the top hits, which showed co-localized signal with TRA-1-60 (Figures S4a-c). Importantly, D8 cells enriched with the candidate probes gave rise to a significantly higher number of TRA-1-60+ colonies (Figures 4a and S4d). Among them, BDD2-C8 displayed the best performance in an alternative reprogramming of MRC5 fibroblasts (Figure S4e). BDD2-C8 also precisely captured changes in reprogramming efficiency upon depletion of the key modulators^15^ (Figure S4f). Single-cell qPCR showed that BDD2-C8+ cells expressed higher levels of epithelial and pluripotent genes, and lower levels of mesenchymal and somatic genes (Figure 4b). To further characterize, we prepared 192 scRNA-Seq libraries for D8 cells stained highly (D8^BDD2-C8+^) and lowly (D8^BDD2-C8-^) for BDD2-C8. In the ensuing RCA analysis, D8^BDD2-C8+^ and D8^BDD2-C8-^ cells clustered close to the D8 G2-3 and G1 respectively, which were substantiated by the similar expression profiles for the subgroup specific genes (Figures 4c-d and S4g-h). GO enriched for D8^BDD2-C8+^ specific genes were related to cell BDD2-C8-cycle, embryo development and stem cell population maintenance, whereas D8^BDD2-C8-^ genes were predominantly represented by epithelial to mesenchymal transition, extracellular matrix organization, and development processes (Figure 4e and Supplementary Table 4). In terms of BDD2-C8-structural complexes, D8^BDD2-C8-^ genes were specifically enriched in the endoplasmic reticulum (ER) lumen and Golgi (Figure S4i). Interestingly, BDD2-C8 were localized in the ER and Golgi (Figure S4j). In addition, higher expression of secretory genes in the D8^BDD2-C8-^ cells implicated its active ER-Golgi secretion pathway (Figure S4k). Depletion of these genes indeed resulted in the retention of BDD2-C8 (Figures S4l-m). This indicates that BDD2-C8 may be actively effluxed from BJ and the non-reprogrammed cells (D8 G1) but retained in the pluripotent cells and intermediate cells with high reprogramming potential, due to the differential ER-Golgi secretion activities.

**Figure 4.**
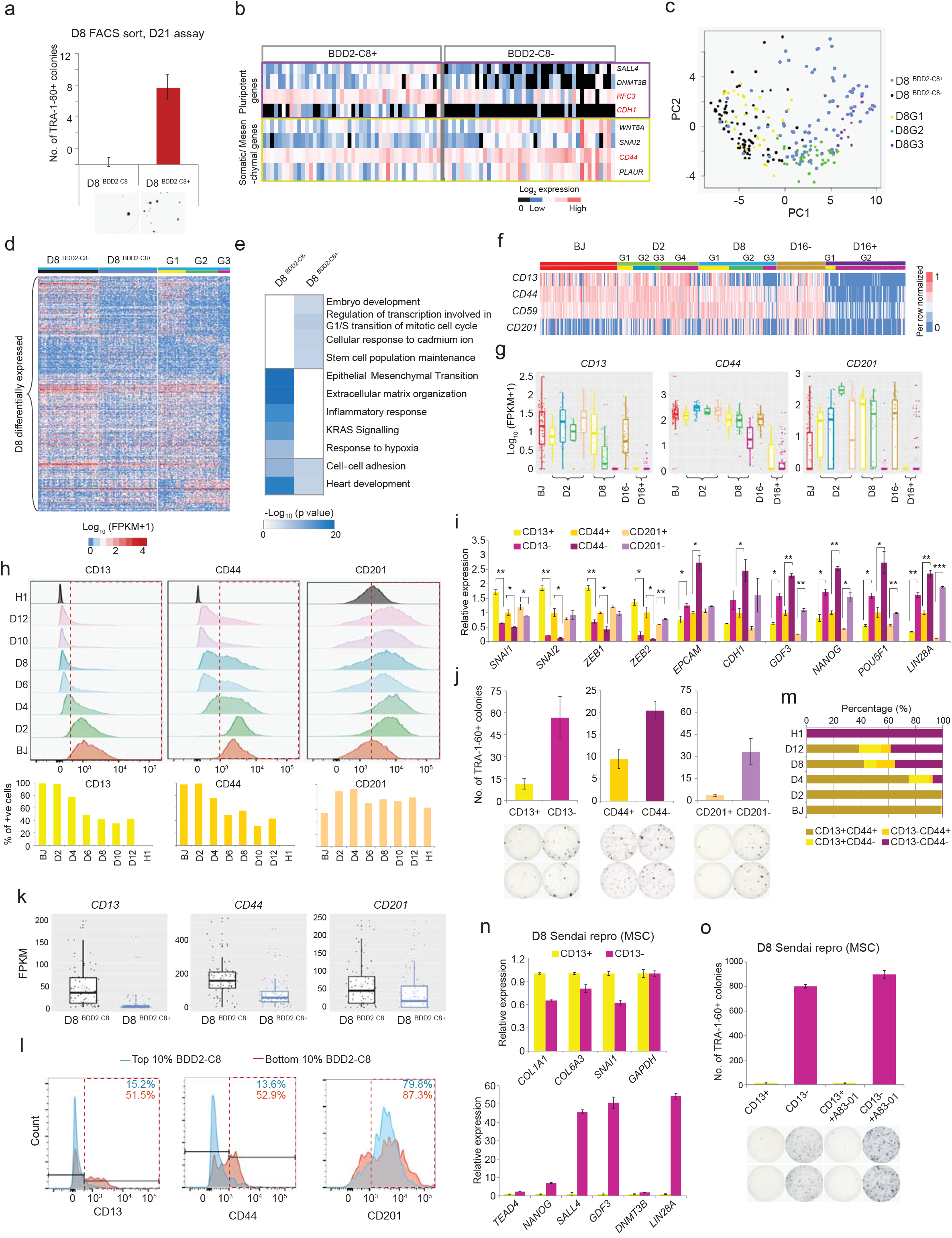
Identification of chemical dyes and surface markers for early-intermediate reprogramming cells. (a) Quantification of TRA-1-60+ colonies yielded from D8 cells sorted with BDD2-C8 (top). Representative images are shown below. n=3; error bar indicates SD. (b) Gene expression in the D8 BDD2-C8+ and BDD2-C8-cells, measured by single cell qRT-PCR. Student t-test (2-tails) was applied (p<0.05). (c) RCA plot of D8, D8^BDD2-C8+^ and D8^BDD2-C8-^ cells. (d) Expression heatmap of the D8 differential genes in D8^BDD2-C8+^ and D8^BDD2-C8-^ cells. Color code on top represents the time points (above) and the subgroups or sorted cells (below). (e) GO terms enriched by the differential genes of D8^BDD2-C8+^ and D8^BDD2-C8-^ cells. (f) Heatmap showing the expression dynamics of the surface markers. Color code on top represents the time point and their respective subgroups. (g) Boxplots showing the surface markers expression across the time points and their respective subgroups. Lines in the box represent the median expression. (h) Stacked histograms (top) showing the fluorescence intensity (X-axis) of the surface markers. Red dotted boxes highlight the positively stained cells. Quantifications are shown below. (i) Barchart exhibiting the relative expression levels in the D8 sorted cells, normalized to that of D8 CD44+ cells. n=2; error bar indicates SD. (j) Quantification of TRA-1-60+ colonies yielded from the D8 sorted cells. Representative images are shown below. n=2; error bar indicates SD. (k) Boxplots showing the surface marker expression in the D8^BDD2-C8+^ and D8^BDD2-C8-^ cells. Lines represent the median expression. (l) Overlaid histograms showing the surface marker staining intensity in the BDD2-C8+ and BDD2-C8-cells. Red dotted boxes highlight the positively stained cells. The numbers on top indicate the percentage of positively stained cells. (m) Barchart showing the distribution of co-staining signals of CD13 and CD44 in the cells of various reprogramming time points and cell lines. (n) Barcharts exhibiting the relative expression of the collagen/ mesenchymal genes (top) and pluripotent genes (bottom) in the D8 CD13 sorted cells. n=2; error bar indicates SD. (o) Quantification of TRA-1-60+ colonies yielded from D8 CD13 sorted cells (top). Representative images are shown below. n=2; error bar indicates SD.

We next identified surface markers with differential expression among D8 subgroups to enrich for the intermediate cells with varying reprogramming potentials (Supplementary Table 4). Shortlisted surface markers displayed higher expression in D8 G1/G2 and D16-cells than D8 G3 and D16+ cells respectively (Figures 4f-g). Majority of them were abundantly enriched in the parental BJ cells, except for *CD201*. Expression dynamics of the surface markers were validated by time-course FACS analysis (Figure 4h). Further, D8 cells stained negatively for the surface markers exhibited lower expression of mesenchymal markers but higher epithelial and pluripotency genes, including the D8 G3 marker *GDF3* (Figure 4i). Noteworthy, negatively stained D8 populations gave rise to more TRA-1-60+ colonies, indicating the capacity of surface markers to isolate early reprogrammed cells with high stemness feature (Figure 4j). We then examined the similarity of cells sorted by BDD2-C8 and the identified surface markers. Indeed, D8^BDD2-C8+^ cells demonstrated lower level of CD13, CD44, and CD201 (Figure 4k-l). Of note, difference in the CD201 protein amount was subtler, which could be due to its inconsistent expression across the D8 subgroups (G1-2 like) with BDD2-C8 (G2-3 like). Of note, co-staining of CD13 and CD44 markers showed extensive overlaps across the time-points (Figures 4m and S4n). Surface markers were verified in an alternative reprogramming of MSCs induced by Sendai viruses (Figure S4o-p). D8 sorted MSC reprogramming cells demonstrated the similar expression trends as BJ reprogramming (Figures 4n and S4q). Importantly, CD13-MSCs resulted in a remarkably higher reprogramming efficiency with or without the aid of TGF-beta inhibition (Figure 4o). Collectively, the fluorescent dye and surface markers established the technological platforms to enrich for the minority of intermediate cells primed for successful reprogramming.

### Refined classification of the intermediate population

To decipher the intermediate reprogramming cells, we prepared 10X libraries on D8 cells sorted with CD13. Majority of the libraries passed the QC thresholds (Figure S5a). Expectedly, libraries showed 2 distinct groups with differential *CD13* expression (Figures 5a and S5b). Notably, majority of the genes that were expressed highly in CD13+ cells were correspondingly expressed in the D8^BDD2-C8-^ cells and vice versa (Figure S5c). The libraries demonstrated 8 clusters. Clusters 5-8 were mainly comprised of CD13+ cells (Figures 5b and S5d). Consistently, CellNet analysis showed that clusters 1-4 correlated with ESCs, whereas the other clusters to the fibroblast state (Figure S5e). RCA corroborated the association of CD13+ cells to the MSCs and Fibroblast lineages (resembling G1) (Figure 5c). Interestingly, among the CD13-cells, cluster 1 showed the highest correlation score to the PSCs, indicating similarity to the D8 G3 cells (Figure 5c). Whereas, cells of cluster 3 and 4 showed resemblance to both the MSCs and ESCs, albeit at a lower significance than cluster 1 in terms of the ESCs (Figure 5c). On the other hand, cells of cluster 2 and 7 exhibited a transitional intermediate profile (Figure 5c). We next used MAGIC^32^ for pairwise comparison between *CD13* and *GDF3*. Interestingly, cells with lower *CD13* levels exhibited higher expression of *GDF3* and *NANOG*, among which cluster 1 and 4 exhibited the highest *GDF3* expression (Figures 5d-e and S5f). Next, to trace the trajectory of the CD13 sub-clusters, we aggregated the CD13 sorted 10X libraries with the other timepoints and performed pseudotemporal analysis. Remarkably, D8 CD13+ and CD13-cells were enriched at the distinct trajectory states (Figure 5f). Cells of cluster 1 and 4 concentrated in the branch shared by D16+ cells. Notably, cells of cluster 3 were found at the unsuccessful reprogramming branch, whereas cluster 2 cells were scattered.

**Figure 5.**
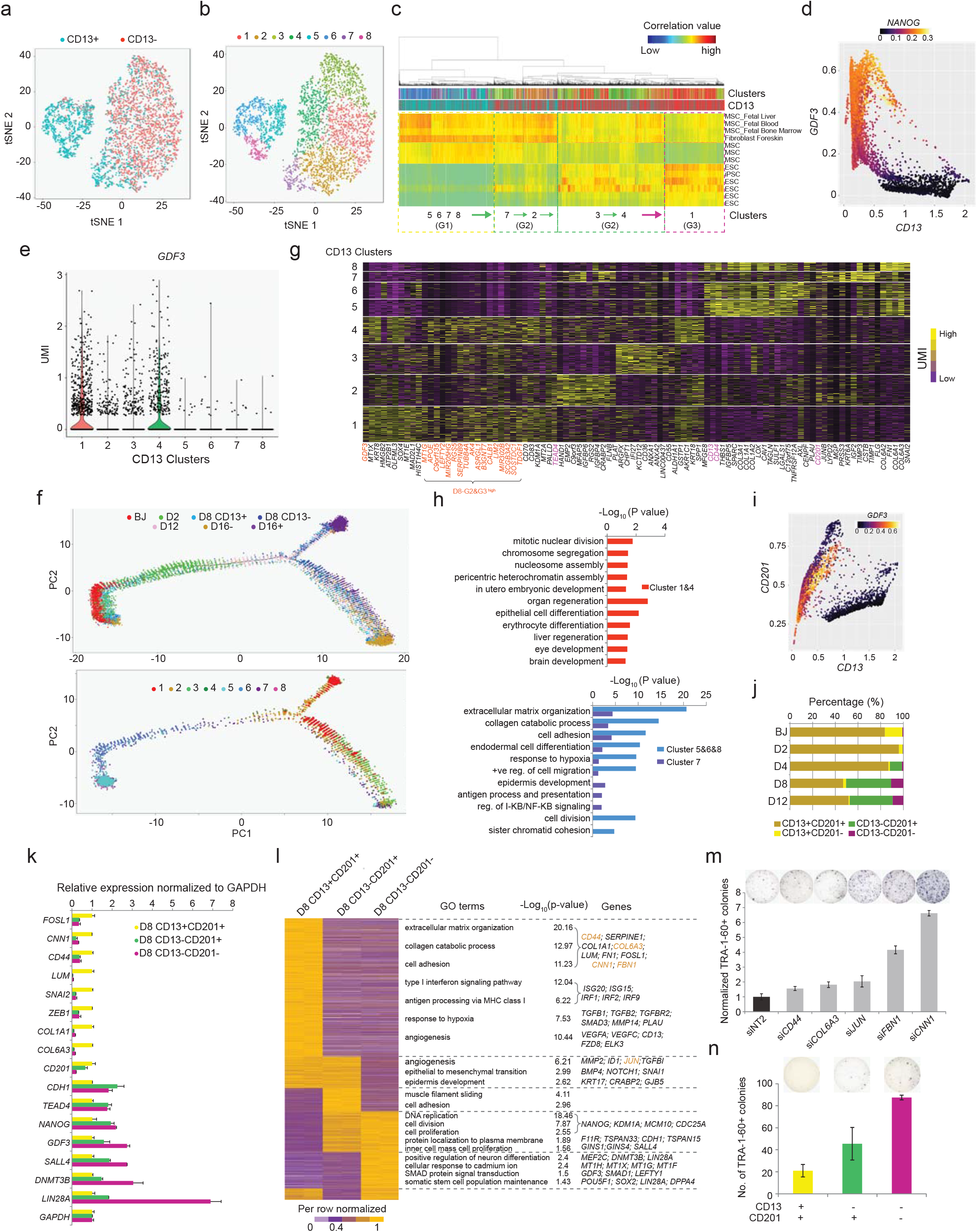
Refined classification and enrichment of early-intermediate reprogramming cells. (a) t-SNE plot generated by 10X libraries of D8 CD13 sorted cells. (b) t-SNE plot showing the clusters among the CD13+ and CD13-cells. (c) RCA heatmap demonstrating the clustering of CD13 sorted cells (column) based on their correlation to the cells of different lineages (row). Color code on top indicates the CD13 antigen profile (below) and clusters (above). (d) MAGIC plot showing the correlative expression of *CD13* and *GDF3* in the D8 CD13-sorted 10x libraries. Color represents the expression level of *NANOG*. (e) Violin plot demonstrating the expression of *GDF3* across the identified clusters. (f) Top: Reprogramming trajectory constructed by the 10X scRNA-Seq libraries of various time points and the D8 CD13 sorted cells. Bottom: Super-imposition of CD13 clusters on the reprogramming trajectory. (g) Heatmap showing the differentially expressed genes of CD13 clusters. Genes highlighted in orange were expressed higher in D8 G2 and G3 than G1. (h) Barcharts showing the GO terms enriched by the genes highly expressed in the indicated CD13 clusters. (i) MAGIC plot showing the correlative expression of *CD13* and *CD201* in the CD13 10X libraries. Color represents the expression level of *GDF3*. (j) Barchart showing the distribution of co-staining signals across the various reprogramming time points and cell lines. (k) Barchart exhibiting the relative gene expression in the D8 dual antibody sorted cells, normalized to that in CD13+CD201+ cells. n=2; error bar indicates SD. (l) Left: Heatmap showing the differentially expressed genes among the D8 dual antibody sorted cells. GO terms (Middle) enriched by the differentially expressed genes, with the enrichment values and the related genes indicated on the right. (m) Bar chart demonstrating the number of normalized TRA-1-60+ colonies (Y-axis) upon the knock-down of genes highly expressed in CD13+ CD201+ or CD13-CD201+ cells, at day 5 of reprogramming. Representative images are shown above. n=3. Error bar indicates SD. (n) Quantification of TRA-1-60+ colonies yielded from the D8 dual antibody sorted cells. Representative images are shown above. n=2; error bar indicates SD.

DGE analysis demonstrated that cluster 7 had unique expression trends among the CD13+ clusters, which associated with epidermis development, and I-KB/NF-KB signaling (Figures 5g-h). On the other hand, cluster 5, 6 and 8 specific genes related to extracellular matrix organization and collagen catabolic process (Figure 5h). cluster 1 and 4 specific genes related to cell division, heterochromatin assembly, embryonic development, organ development of multiple lineages, and MAPK cascade, suggesting that reprogramming cells of cluster 1 and 4 might adopt an open chromatin structure, especially at genes crucial for multi-lineage development and differentiation. Mathematical imputation showed an extensive correlation between *CD13* and *CD44* (Figures S5g-h). Interestingly, we detected a group of cells which were low in *CD13* but high in *CD201*, representing the D8 G2-like intermediate cells belonging mostly to cluster 2 and 7 (Figures 5i, and S5i). To substantiate their existence, we performed time-course co-staining using CD13 and CD201 antibodies. A significant number of intermediate cells were CD13-CD201+, and their proportion increased as reprogramming advanced (Figures 5j and S5j). Furthermore, CD13-CD201+ cells corresponded to higher BDD2-C8 staining signals than CD13+CD201+ cells (Figure S5k). Hence, we hypothesized that dual sorting with CD13 and CD201 markers would allow us to enrich for successfully reprogrammed early-intermediary cells with higher purity.

We thus categorized D8 cells into double negative (CD13-CD201-), double positive (CD13+CD201+) and intermediate (CD13-CD201+) cells, which were then subjected to gene measurement (Supplementary Table 5). CD13+CD201+ cells expressed fibroblast and mesenchymal genes and genes associated with extracellular matrix and cell adhesion (Figures 5k-l). On the contrary, CD13-CD201-cells expressed genes related to pluripotency, epithelial lineage, cell division, neuronal differentiation, and stem cell population maintenance. CD13-CD201+ cells exhibited an intermediate transcription profile (Figures 5k-l). In addition, genes highly expressed in CD13+CD201+ cells were mostly enriched in the D16-but not D16+ cells, whereas the opposite was observed for CD13-CD201-specific genes (Figure S5l). Notably, depletion of genes highly expressed in the CD13+CD201+ population resulted in more reprogrammed colonies (Figure 5m). Importantly, enrichment of CD13-CD201-population gave rise to the highest reprogramming efficiency, in comparison with the CD13-CD201+ (intermediate) and CD13+CD201+ cells (lowest) (Figure 5n). We validated the presence of these distinct D8 populations in an alternative reprogramming of BJ cells using Sendai viruses (Figures S5m-o). Altogether, concurrent use of CD13 and CD201 antibodies enable us to dissect the precise populations differentially poised for successful reprogramming.

### Regulatory networks of Transcription Factors in cellular reprogramming

Transcription Factors (TFs) define the cell-selective regulatory network underlying the cellular identity and function^33^. However, the stage-specific core regulatory networks of the human cellular reprogramming remain elusive. To this end, we performed DGE analysis for TFs across the pseudotemporal states, which were then categorized as Early Silenced, Late Silenced, Transient, Early Expressed and Late Expressed (Figures 6a and S6a, and Supplementary Table 6). Notably, many TFs exhibited similar expression trends in the D8 RCA subgroups and the D8 sorted libraries. Specifically, Early Silenced TFs (e.g. *FOSL1*, *CREB3L1*, *AHRR*, *DRAP1* and *ELL2*) showed higher expression in the D8 BDD2-C8- and CD13+CD201+ populations, whereas Late Silenced TFs exhibited the opposite trends (Figures 6a and S6b). Majority of Transient TFs exhibited higher expression in CD13+CD201+ and CD13-CD201+ populations, including lineage associated factors namely *HAND1* (mesoderm), *ASXL3* and *NEUROG2* (neuroectoderm) (Figures 6a and S6c). Contrastingly, Early Expressed TFs adopted higher expression in the cells of CD13-CD201- and CD13-CD201+ (Figures 6a and S6d). CD13-CD201-cells expressed the highest level of Late Expressed TFs, such as *PRDM14*, *DNMT3B* and *LHX6*.

**Figure 6.**
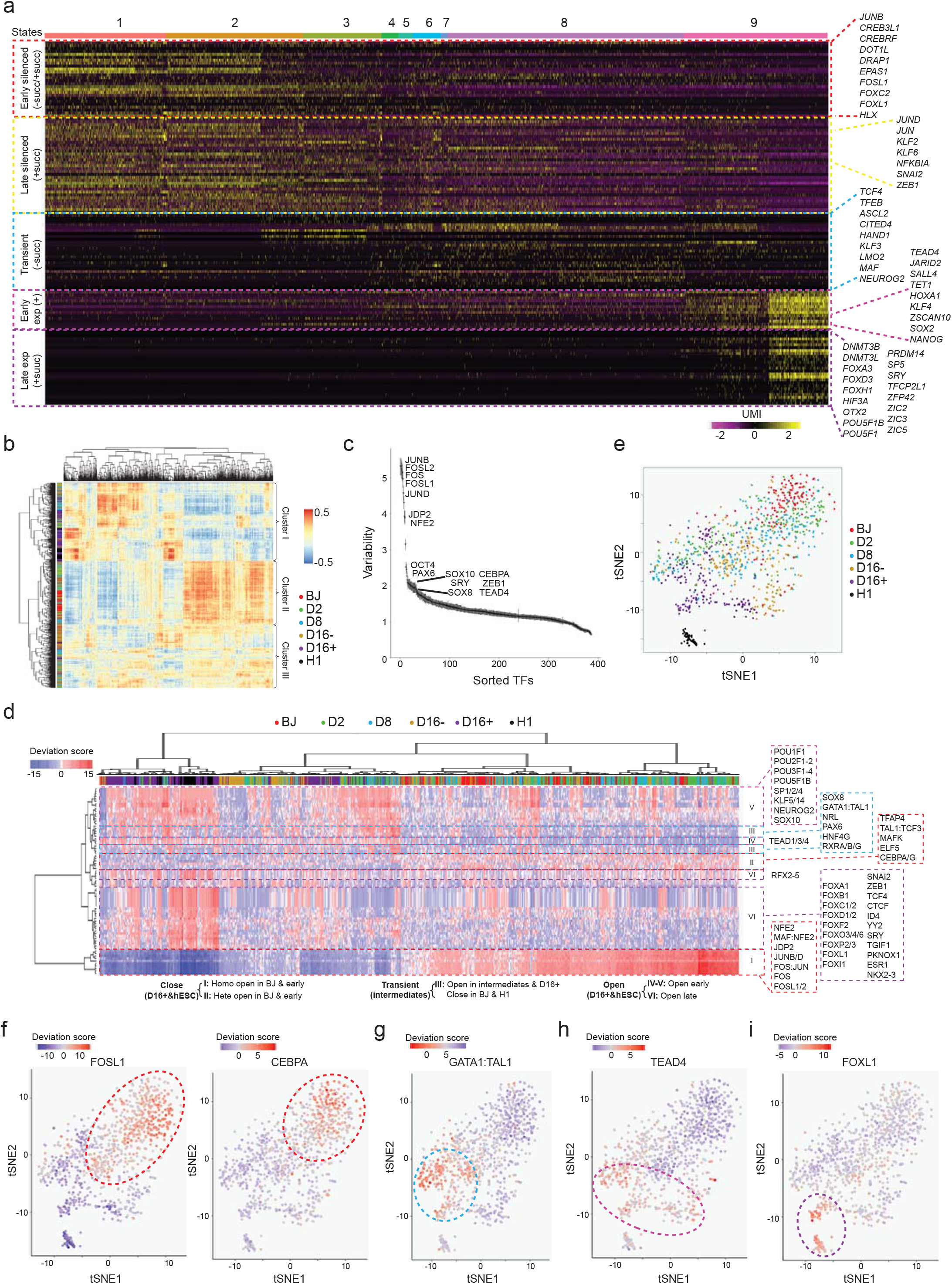
Transcription factors critical for reprogramming. (a) Heatmap showing the expression dynamics of the transcription factors across the pseudotime states. Color represents the expression level, ranging from purple (low) to yellow (high). Color code on top represents the pseudotime states. Transcription factors were classified to 5 categories according to the states when they were silenced or expressed. The representative TFs of each category are listed on the right. (b) Correlation heatmap of scATAC-Seq libraries based on the calculated JASPAR motifs deviations in the HARs peaks. Color represents the correlation value, ranging from blue (no) to red (high). Side color bar indicates the time-point to which each scATAC-Seq library belongs. (c) Plot indicating the significantly variable motifs in terms of accessibility from the scATAC-Seq libraries. Y-axis represents the variability score assigned to each JASPAR motif whereas the X-axis represents the motif rank. (d) Heatmap of scATAC-Seq libraries based on the deviation scores of the significantly variable JASPAR motif. Color indicates the accessibility level, ranging from dark blue (no enrichment) to dark red (high enrichment). Color code on top represents the time points. Motifs were classified to 3 major types according to the dynamics of accessibility across the time points. (e) t-SNE plot of scATAC-Seq libraries based on the deviation scores of JASPAR motifs. (f-i) Super-imposition of motif enrichment scores for Close motif-FOSL1 and CEBPA (f), Transient motif-GATA1:TAL1 (g), Open motif: Early Open-TEAD4 (h); Late Open-FOXL1 (i) on the scATAC-Seq tSNE plot. Color indicates the accessibility level and ranges from dark blue (no enrichment) to dark red (high enrichment).

To investigate how the regulatory TFs accessed its genomic targets, we then analyzed time-course scATAC-Seq libraries of reprogramming cells (Figure 1a). Both batches of libraries showed similar promoter accessibility profiles (Figure S6e). We also observed diminished accessibility on the fibroblast-specific DHS and increased accessibility on the pluripotency-specific DHS as reprogramming progressed (Figure S6f). Next, we utilized chromVAR^34^ to identify the TFs determining the variable epigenomes accessibility. Correlation of scATAC-Seq libraries among themselves resulted in three major clusters. The early cells consisting mostly of BJ and D2 cells clustered together (Cluster II), while D8, D16+ and H1 cells shared similar accessibility profile (Cluster I). A third cluster composed mainly of D8 and D16-cells (Cluster III) (Figure 6b). of note, FOSL1 and its partners, CEBPA, ZEB1, PAX6, SOX8, SOX10, POU5F1 and TEAD4 were found to be the TFs contributing to the reprogramming heterogeneity (Figure 6c and Supplementary Table 6). TFs were then categorized to open starting from the intermediate stage (Early open) or the late stage (Late open), open transiently at the intermediate stage (Transient open), and close starting from the intermediate stages of reprogramming (Close) which were either homogenously or heterogeneously open in the early cells (Figure 6d and 6e). Particularly, Close TFs belonged mostly to the FOS-JUN-AP1 complex, such as FOSL1 and JDP2 (Figures 6f and S6g). Intriguingly, motifs of lineage specifiers (Mesendoderm: GATA1, SOX8 and HNF4G; Ectoderm: PAX6, NRL and RXR) were found to be accessible only in the intermediate reprogramming cells, but not the H1 hESCs (Figures 6d, 6g and S6h). This observation corroborated with the model of counteracting lineage specification networks underlying the induction of pluripotency^35,36^. On the contrary, Early Open TFs (e.g. TEAD4 and POU5F1) exhibited accessibility starting from the D8 reprogramming cells, whilst Late Open TFs (e.g. FOXL1, TCF4, and YY2) demonstrated accessibility in D16+ cells and ESCs exclusively (Figures 6d, 6h-i, S6i-k). Of note, TFs of FOX family and YY1 were previously reported to be important for reprogramming^37–39^. Independently, we used SCENIC analysis to infer the interaction between TFs and their target genes. Corroborating with our earlier results, Silenced and Close TFs showed decrease in their regulon activity as reprogramming progressed, whereas majority of Expressed and Open TFs displayed the opposite dynamics (Figure S6l). Likewise, regulon activity of the Transient TFs was observed only in the intermediate cells. Together, these data represent the compendium of TFs which regulate the networks of downstream key modulators in the cellular reprogramming process.

### Identification of key regulators for the intermediate stage of reprogramming

In order to deduce TFs essential for the intermediate cells to acquire pluripotency, we further sub-clustered the D8 scATAC-Seq libraries (Figures S7a-b). Intriguingly, the most variable motifs of D8 cells belonged to the FOS-JUN-AP1 and TEAD families (Figure 7a and Supplementary Table 7). Strikingly, D8 cells were either accessible for FOSL1-JUN-AP1 or TEAD4 motif (Figures 7b-c). Of note, *FOSL1* and *TEAD4* displayed a contrasting expression pattern and regulon activity during reprogramming (Figures 7d-g). In D8, *FOSL1* and *TEAD4* were exclusively expressed in the CD13+ and CD13-cells (Figure 7h).

**Figure 7.**
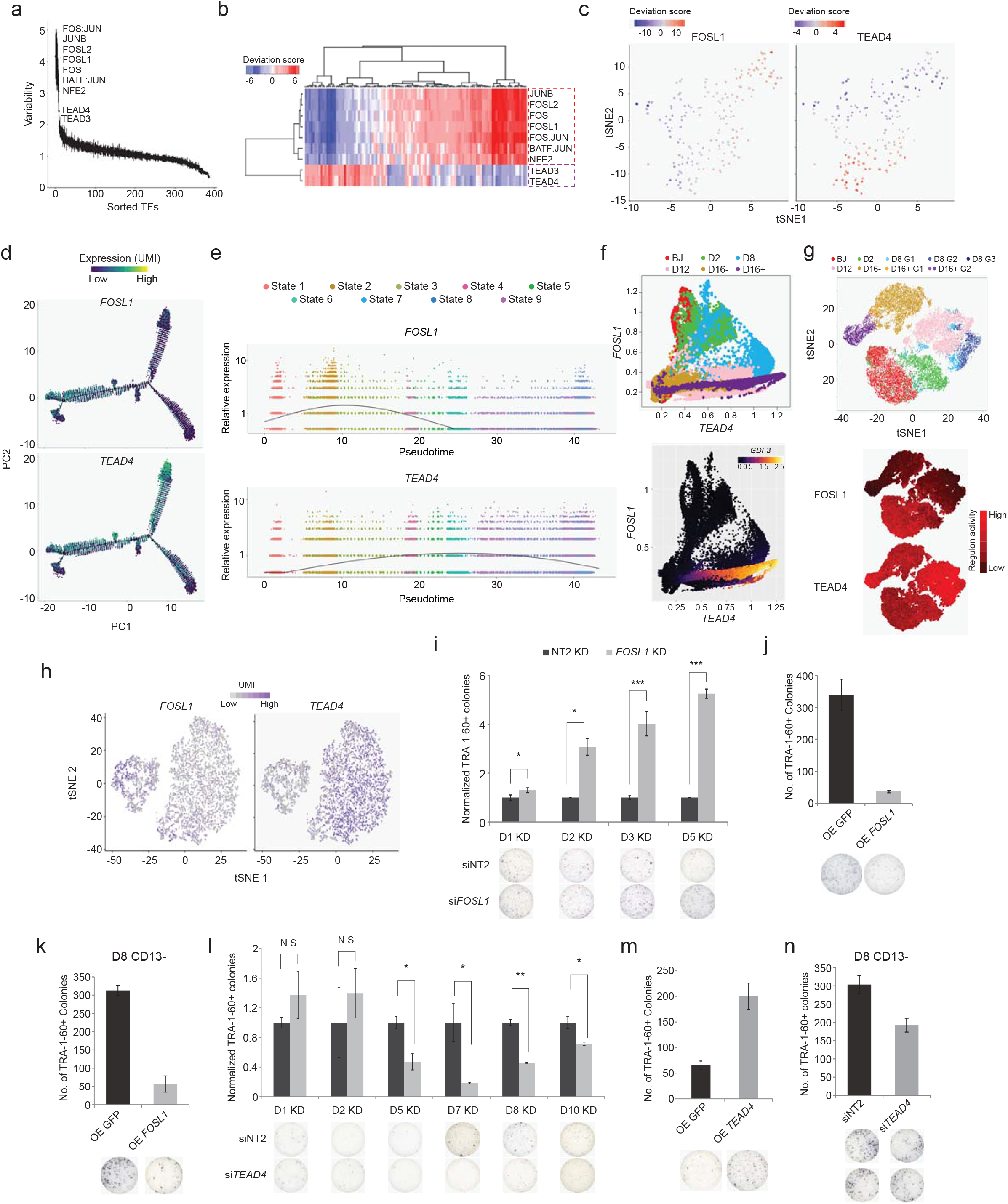
Transcription factors contributing to the heterogeneity in chromatin accessibility of the intermediate reprogramming cells. (a) Plot indicating the significantly variable motifs in terms of accessibility among D8 cells. Y-axis represents the variability score assigned to each JASPAR motif whereas X-axis represents the motif rank. (b) Heatmap showing clustering of D8 scATAC-Seq libraries based on the deviations scores of the significantly variable JASPAR motif. (c) Super-imposition of motif enrichment scores for FOSL1 and TEAD4 on the D8 scATAC-Seq tSNE plot. (d) Super-imposition of *FOSL1* and *TEAD4* expression on the reprogramming trajectory displayed in Figure 3a. (e) Dotplots indicating the expression of *FOSL1* and *TEAD4* along the pseudotime. Smooth lines represent the mean expression level at each pseudotime, regardless of the state. (f) Top: MAGIC plot showing correlative expression of *FOSL1* and *TEAD4* in the 10X libraries. Bottom: Super-imposition of *GDF3* expression on the magic plot. (g) Top: tSNE plot based on the regulon activity matrix generated by SCENIC. Bottom: Activity of FOSL1 and TEAD4 regulons across the time-points. (h) Super-imposition of *FOSL1* and *TEAD4* expression on the D8 CD13 sorted tSNE plot. (i) Bar chart demonstrating the number of normalized TRA-1-60+ colonies upon the knock-down of *FOSL1* at the indicated time-points of reprogramming. Representative images are shown below, n=3. Error bar indicates SD. (j) Bar chart demonstrating the quantification of TRA-1-60+ colonies upon the overexpression of *FOSL1* at D5. Representative images are shown below, n=3. Error bar indicates SD. (k) Bar chart demonstrating the quantification of TRA-1-60+ colonies yielded from D8 CD13-cells with *FOSL1* overexpression. Representative images are shown below, n=3. Error bar indicates SD. (l) Bar chart demonstrating the number of normalized TRA-1-60+ colonies upon the knockdown of *TEAD4* at the indicated time-points of reprogramming. Representative images are shown below, n=2. Error bar indicates SD. (m) Bar chart demonstrating the quantification of TRA-1-60+ colonies upon the overexpression of *TEAD4*. Representative images are shown below, n=3. Error bar indicates SD. (n) Bar chart demonstrating the number of TRA-1-60+ colonies yielded from D8 CD13-cells with *TEAD4* depletion. Representative images are shown below, n=3. Error bar indicates SD.

Given its expression and motif accessibility at the early stage of reprogramming, FOSL1 was hypothesized to act as a roadblock. To verify, reprogramming efficiency was assessed upon the depletion of *FOSL1* by siRNA, which showed no effect on cell proliferation and lasted for about 4 days (Figures S7c-e). Remarkably, depletion of *FOSL1* at day 1, 2, 3, or 5 post-OSKM induction resulted in an increased reprogramming efficiency (Figure 7i). Specificity of the *FOSL1* siRNA construct was validated using a mutant construct which showed no effect (Figures S7f-g). Depletion of *FOSL1* in an alternative Sendai virus reprogramming reproduced the phenotypic change (Figures S7h-i). Conversely, ectopic expression induced a drastic reduction in the number of reprogrammed colonies (Figure 7j). Noteworthy, high reprogramming propensity of CD13-cells was negated by the elevated *FOSL1* level (Figure 7k).

Contrastingly, we posit that TEAD4 could serve as an effector. Indeed, depletion of *TEAD4* resulted in the reduced number of reprogrammed colonies (Figure 7l). Specificity of si*TEAD4* was affirmed by the mutant construct with no phenotypic change (Figures S7j-k). Knock-down effect of si*TEAD4* lasted for around 5 days (Figure S7l). Interestingly, when knock-down was performed on D5 cells, *TEAD4* expression did not restore, which could be due to the perpetuations of the non-reprogrammed state introduced by si*TEAD4* (Figure S7m). Notably, depletion of *TEAD4* only from day 5 onwards induced the decreased reprogramming efficiency (Figure 7l). This established the vital role of TEAD4 at the intermediate-late stages of reprogramming. In contrast, overexpression of *TEAD4* resulted in a significant increase in the number of reprogrammed colonies (Figure 7m). Importantly, depletion of *TEAD4* revoked the reprogramming potential of D8 CD13-cells (Figure 7n). Collectively, these analyses illustrate that the pivot from FOSL1 to TEAD4-directed regulatory network is essential for successful reprogramming.

### Mechanistic roles of FOSL1 and TEAD4 in cellular reprogramming

To find the direct targets of FOSL1, we prepared ChIP-Seq libraries for D8 cells, which demonstrated the expected genomic distribution profile and enriched with FOSL1 motif and its regulatory partners (Figures S8a-c). We also investigated the genomic binding profile of TEAD4 in D8 cells and CD13 sorted D8 cells (Figures S8d-e). Majority of the D8 TEAD4 bound sites were shared by both CD13+ and CD13-cells (Figures S8f). 1.7-fold higher numbers of TEAD4 bound loci were detected in the CD13-than CD13+ cells (18550 vs. 10986 sites). In addition, CD13-specific sites showed greater enrichment of TEAD4 motif and exclusive enrichment of pluripotency associated OCT4-SOX2-TCF-NANOG motif, indicating the differential regulatory role of TEAD4 (Figures S8g).

Consistent with their contrasting roles, majority of loci highly enriched with FOSL1 were distinctive from that with TEAD4 (Figure 8a). We next clustered BJ, D16- and D16+ scATAC-Seq libraries, based on the accessibility of FOSL1 and TEAD4 bound sites (Figure 8b). Notably, FOSL1 specific sites exhibited higher accessibility in BJ and D16-cells, whereas CD13-specific and CD13 common TEAD4 bound sites were mostly accessible in D16+ cells (Figures 8b-c and S8h-i). these differential accessible sites were annotated as the functional FOSL1 and TEAD4 targets (Supplementary Table 8). Similar to the motif enrichment pattern, most of the D8 cells were either accessible for FOSL1 (“FOSL1 ChIP only” cells) or CD13-TEAD4 bound sites (“TEAD4 ChIP only” cells) (Figures 8d-e and S8k-l). We next asked if their binding has any consequence to the transcription of target genes. To this end, we analyzed the expression of functional FOSL1 and TEAD4 targets across the D8 CD13-CD201 sorted cells. Remarkably, majority of the D8 FOSL1 targets (e.g. *TGFBR2*, *SMAD3*, and *COL5/7/21A1*) were expressed highly in the CD13+CD201+ cells (Figure 8f). In contrast, D8 CD13-CD201-cells demonstrated high expression of the TEAD4 targets, including key pluripotent genes such as *DNMT3B*, *LIN28A*, *SOX2*, and *PODXL* which codes for TRA-1-60 (Figure 8g). This was further substantiated at single-cell resolution using the coupled NMF analysis of D8 scRNA-Seq and scATAC-Seq libraries, by which regulatory connectivity between the accessible chromatin regions and expression of target genes were established (Figure S8m). Notably, cluster 1 composed of “TEAD4 ChIP only” cells (scATAC-Seq) and D8 RCA G3 cells (scRNA-Seq), whilst cluster 2 comprised of “FOSL1 ChIP only” cells (scATAC-Seq) and D8 RCA G1 (scRNA-Seq) cells. Differential analysis demonstrated that TEAD4 and its targets, such as TET1 and CDH1, were highly accessible and expressed in the cluster 2 cells, whereas FOSL1 and its targets, such as MMP2 and SMAD3, displayed the opposite trends (Figures S8m-n). Besides, interactome analysis revealed that cluster 1 genes associated with splicing process, whereas cluster 2 genes related to ER-Golgi transport and extracellular matrix organization, including FOSL1 targets *MMP2* and several collagen genes (Figure S8o). More importantly, downstream targets of FOSL1 and TEAD4 were themselves modulators of reprogramming. For instance, knock-down of FOSL1 bound genes, *MMP2* and *SMAD3*, resulted in higher numbers of the reprogrammed colonies (Figure 8h). Conversely, depletion of TEAD4 targets, *PRDM14* and *SOX2*, showed the opposite effect (Figure 8i).

**Figure 8.**
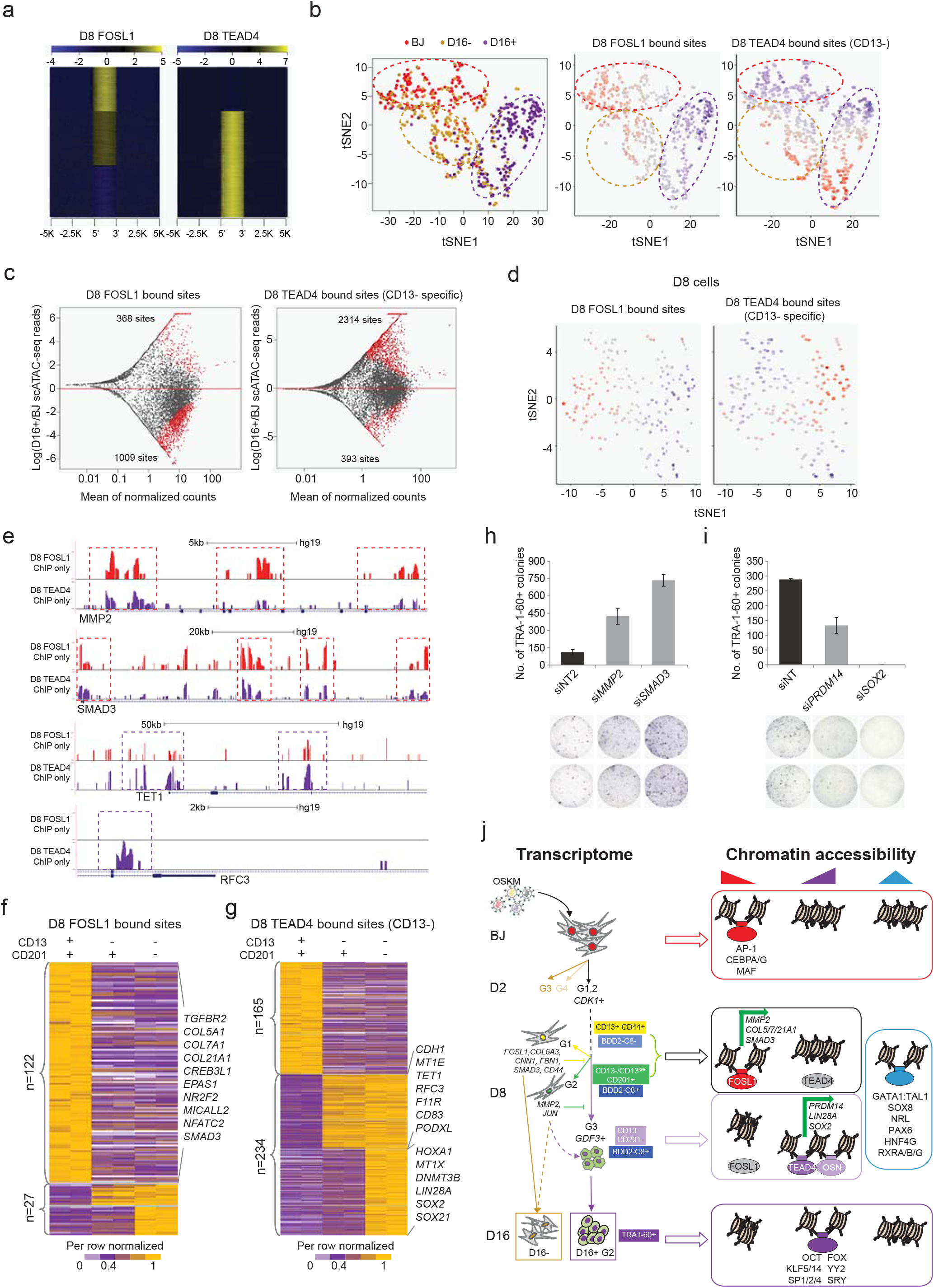
Mechanistic role of FOSL1 and TEAD4 during reprogramming. (a) Heatmaps exhibiting the D8 FOSL1 and D8 TEAD4 ChIP-Seq enrichment over merged binding loci of both these factors. (b) Left: t-SNE clustering of BJ, D16+, and D16-scATAC-Seq libraries based on the deviation scores of common and specific sites identified from D8 FOSL1 ChIP-seq, D8 CD13-TEAD4 ChIP-seq, and D8 CD13+ TEAD4 ChIP-seq. Right: Super-imposition of deviation score for FOSL1 specific bound sites and TEAD4 CD13-specific bound sites on the scATAC-Seq t-SNE plot. Color indicates the accessibility level. (c) MA plots of scATAC-Seq revealing the differentially accessible FOSL1 bound sites (left) and CD13-specific TEAD4 bound sites (right) between BJ and D16+ cells. X-axis denotes the mean of normalized counts in BJ cells. Red dots denote the sites with significant accessibility changes. (d) Super-imposition of enrichment scores for FOSL1 specific bound sites (left) and CD13-TEAD4 specific bound sites (right) on the D8 scATAC-Seq tSNE plot. Color indicates the accessibility leve. (e) UCSC screenshots illustrating the chromatin accessibility of FOSL1bound genes (MMP2 and SMAD3) and TEAD4 CD13-bound sites (TET1 and RFC3) in D8 “FOSL1 ChIP only” and D8 “TEAD4 ChIP only” cells. The differentially accessible sites with highlighted with dotted rectangles. (f-g) Expression heatmap of the differentially accessible genes bound by FOSL1 (f) in D8 and TEAD4 in D8 CD13-cells (g), among the D8 dual antibody sorted cells. (h-i) Bar chart demonstrating the number of TRA-1-60+ colonies upon knock-down of FOSL1 targets (h) and TEAD4 targets (i) at D5. Representative images are shown below. n=3. Error bar indicates SD. (j) Proposed model of the study.

Taken together, we present the single-cell roadmap of the human cellular reprogramming process, which reveals the diverse cell-fate trajectory of individual reprogramming cells (Figure 8j). D2 cells consist of four subgroups, out of which *CDK1+* cells showed greater propensity for reprogramming. Among the three subgroups of D8 cells, G1 represents the unsuccessful reprogramming cells, whereas G3 cells with *GDF3* expression are highly primed for successful reprogramming. G2 is the intermediary group between G1 and G3. In-house developed fluorescent probe (BDD2-C8) and the identified surface markers (CD13, CD44 and CD201), enables the segregation of the heterogeneous population based on their reprogramming potential. Moreover, TFs analysis reveals the stage-specific regulatory networks of reprogramming. Importantly, we describe the crucial switch from a FOSL1 to a TEAD4-centric expression which collectively regulate genomic accessibility, cell-lineage transcription program, and network of functional downstream modulators favoring the acquisition of the pluripotent state.

## DISCUSSION

### Heterogeneity of Cellular Reprogramming

Due to the inherent heterogeneity, Single-cell NGS techniques are more suited to characterize the molecular and epigenetics signatures of reprogramming cells with diverse trajectories. Several studies described the mouse reprogramming process at single-cell resolution^40–42^. However, there have been reports for extensive differences between the mouse and human reprogramming in terms of kinetics, modulators involved, and the molecular nature of the generated iPSCs. In addition, this is the first study presenting a comprehensive human reprogramming roadmap by integrating the transcriptomic changes and the alteration of accessible epigenetic regions at single-cell resolution.

Our analyses also identify an early reprogramming marker *GDF3*, which marks the intermediate cells primed for successful reprogramming. *GDF3* was previously shown to be expressed in pluripotent stem cells and played a role in regulating cell fate via BMP signaling inhibition^43^. We also develop a fluorescent probe, BDD2-C8, and identify a panel of cell surface markers, namely CD13, CD44 and CD201, to distinguish and characterize the diverse subpopulations of the intermediate reprogramming cells. Interestingly, CD44 sorting was previously reported as a mean to isolate reprogrammed cells in the mouse system^44^. Noteworthy, combinatorial use of the surface markers enables a more refined segregation. The toolkits to decipher the intermediate cells with different stemness capacity, will help deepen our understanding of the mechanisms of reprogramming process. Additionally, the ability to enrich for early reprogramming cells will help increase the success rate of iPSC generation from the cell lines or patient-derived primary cells which are refractory to reprogramming.

### Transcription Factors Contribute to the Heterogeneity of Cellular Reprogramming

Facilitated by the integrative analysis of transcriptomic and chromatin accessibility profiles, TFs contributing to the heterogeneous reprogramming trajectories are identified. Accessible regions with the motifs of FOS-JUN-AP1 and CEBPA are rapidly closed upon induction, which were reported to act as repressors in mouse reprogramming^45,46^ (Figure 6). An earlier report showed that FOSL1 lost many of its binding as early as day 2 of mouse reprogramming^46^. Our study reveals that, in the human system, FOSL1 regulates myriad of genes which serve as barriers of reprogramming, including *MMP2*, *SMAD3*, *TGFBR2*, and collagen genes. Strikingly, chromatin regions with the motifs of lineage TFs are open in the intermediate cells, which might be due to the induction of ME and ECT lineages driven by POU5F1 and SOX2^47^. This is further supported by the replacement of POU5F1 and SOX2 with the ME and ECT lineage specifiers, for both mouse and human reprogramming^35,36^. We unravel for the first time the transitory epigenetic accessibility directed by the lineage TFs, which contribute extensively to the diverse cell fate potentials observed during cellular reprogramming.

## ACKNOWLEDGMENTS

We are grateful for Ahad Khalilnezhad for technical assistance and Lin Yang and Samantha Seah for helpful discussion. H.L. is supported by Glenn Foundation for Medical Research, Mayo Clinic Center for Biomedical Discovery and Mayo Clinic Cancer Center. Y-H.L. is supported by the [NRF Investigatorship award-NRFI2018-02] and [JCO Development Programme Grant – 1534n00153] grants. Y-H.L. and L-F.Z. are supported by the Singapore National Research Foundation under its Cooperative Basic Research Grant administered by the Singapore Ministry of Health’s National Medical Research Council [NMRC/CBRG/0092/2015]. We are grateful to the Biomedical Research Council, Agency for Science, Technology and Research, Singapore for research funding.

## AUTHOR CONTRIBUTIONS

Contribution: Q.R.X, C.E.F designed and performed research, analyzed data, and wrote the paper; P.G, Y.S.C, C.X.D.T, T.W designed and conducted research; N.Y.K, S.S, Y.T.C, J.X, J.C, H.L, L.F.Z analyzed data; and Y.H.L. designed research, analyzed data, and wrote the paper.

## AUTHOR INFORMATION

The authors declare no competing financial interests.

## REFERENCE

1. Loh, Y.-H. et al. Reprogramming of T Cells from Human Peripheral Blood. Cell Stem Cell 7, 15–19 (2010).

2. Park, I. H. et al. Reprogramming of human somatic cells to pluripotency with defined factors. Nature 451, 141–146 (2008).

3. Takahashi, K. et al. Induction of Pluripotent Stem Cells from Adult Human Fibroblasts by Defined Factors. Cell 131, 861–872 (2007).

4. Takahashi, K. & Yamanaka, S. Induction of Pluripotent Stem Cells from Mouse Embryonic and Adult Fibroblast Cultures by Defined Factors. Cell 126, 663–676 (2006).

5. Tan, H.-K. et al. Human Finger-Prick Induced Pluripotent Stem Cells Facilitate the Development of Stem Cell Banking. Stem Cells Transl. Med. 3, 586–598 (2014).

6. Yu, J. et al. Induced pluripotent stem cell lines derived from human somatic cells. Science 318, 1917–20 (2007).

7. Seah, Y. F. S., El Farran, C. A., Warrier, T., Xu, J. & Loh, Y. H. Induced pluripotency and gene editing in disease modelling: Perspectives and challenges. Int. J. Mol. Sci. 16, 28614–28634 (2015).

8. Cheow, L. F. et al. Single-cell multimodal profiling reveals cellular epigenetic heterogeneity. Nat. Methods 13, 833–836 (2016).

9. Polo, J. M. et al. A molecular roadmap of reprogramming somatic cells into iPS cells. Cell 151, 1617–1632 (2012).

10. Stadtfeld, M. & Hochedlinger, K. Induced pluripotency: History, mechanisms, and applications. Genes and Development 24, 2239–2263 (2010).

11. Maherali, N. et al. A High-Efficiency System for the Generation and Study of Human Induced Pluripotent Stem Cells. Cell Stem Cell (2008). doi:10.1016/j.stem.2008.08.003

12. Merkl, C. et al. Efficient Generation of Rat Induced Pluripotent Stem Cells Using a Non-Viral Inducible Vector. PLoS One 8, (2013).

13. Borkent, M. et al. A Serial shRNA Screen for Roadblocks to Reprogramming Identifies the Protein Modifier SUMO2. Stem Cell Reports 6, 704–716 (2016).

14. Qin, H. et al. Systematic identification of barriers to human Ipsc generation. Cell 158, 449–461 (2014).

15. Toh, C. X. D. et al. RNAi Reveals Phase-Specific Global Regulators of Human Somatic Cell Reprogramming. Cell Rep. 15, 2597–2607 (2016).

16. Yang, C. S., Chang, K. Y. & Rana, T. M. Genome-wide Functional Analysis Reveals Factors Needed at the Transition Steps of Induced Reprogramming. Cell Rep. 8, 327–337 (2014).

17. Fang, H. T. et al. Global H3.3 dynamic deposition defines its bimodal role in cell fate transition. Nat. Commun. 9, (2018).

18. Gawad, C., Koh, W. & Quake, S. R. Single-cell genome sequencing: Current state of the science. Nature Reviews Genetics 17, 175–188 (2016).

19. Buenrostro, J. D. et al. Single-cell chromatin accessibility reveals principles of regulatory variation. Nature 523, 486–490 (2015).

20. Islam, S. et al. Characterization of the single-cell transcriptional landscape by highly multiplex RNA-seq. Genome Res. 21, 1160–1167 (2011).

21. Ramsköld, D. et al. Full-length mRNA-Seq from single-cell levels of RNA and individual circulating tumor cells. Nat. Biotechnol. 30, 777–782 (2012).

22. Cahan, P. et al. CellNet: Network biology applied to stem cell engineering. Cell 158, 903–915 (2014).

23. Li, H. et al. Reference component analysis of single-cell trans criptomes elucidates cellular heterogeneity in human colorectal tumors. Nat. Genet. 49, 708–718 (2017).

24. Trapnell, C. et al. The dynamics and regulators of cell fate decisions are revealed by pseudotemporal ordering of single cells. Nat. Biotechnol. 32, 381–386 (2014).

25. Qiu, X. et al. Reversed graph embedding resolves complex single-cell trajectories. Nat. Methods 14, 979–982 (2017).

26. Tsubouchi, T. et al. DNA synthesis is required for reprogramming mediated by stem cell fusion. Cell 152, 873–883 (2013).

27. Lu, Y. et al. Alternative splicing of MBD2 supports self-renewal in human pluripotent stem cells. Cell Stem Cell (2014). doi:10.1016/j.stem.2014.04.002

28. Jeong, Y. M. et al. CDy6, a Photostable Probe for Long-Term Real-Time Visualization of Mitosis and Proliferating Cells. Chem. Biol. (2015). doi:10.1016/j.chembiol.2014.11.018

29. Yun, S.-W. et al. Neural stem cell specific fluorescent chemical probe binding to FABP7. Proc. Natl. Acad. Sci. (2012). doi:10.1073/pnas.1200817109

30. Tan, F., Qian, C., Tang, K., Abd-Allah, S. M. & Jing, N. Inhibition of transforming growth factor β (TGF-β) signaling can substitute for Oct4 protein in reprogramming and maintain pluripotency. J. Biol. Chem. (2015). doi:10.1074/jbc.M114.609016

31. Ruetz, T. et al. Constitutively Active SMAD2/3 Are Broad-Scope Potentiators of Transcription-Factor-Mediated Cellular Reprogramming. Cell Stem Cell (2017). doi:10.1016/j.stem.2017.10.013

32. van Dijk, D. et al. Recovering Gene Interactions from Single-Cell Data Using Data Diffusion. Cell (2018). doi:10.1016/j.cell.2018.05.061

33. Neph, S. et al. Circuitry and dynamics of human transcription factor regulatory networks. Cell (2012). doi:10.1016/j.cell.2012.04.040

34. Schep, A. N., Wu, B., Buenrostro, J. D. & Greenleaf, W. J. ChromVAR: Inferring transcription-factor-associated accessibility from single-cell epigenomic data. Nat. Methods 14, 975–978 (2017).

35. Shu, J. et al. XInduction of pluripotency in mouse somatic cells with lineage specifiers. Cell (2013). doi:10.1016/j.cell.2013.05.001

36. Montserrat, N. et al. Reprogramming of human fibroblasts to pluripotency with lineage specifiers. Cell Stem Cell (2013). doi:10.1016/j.stem.2013.06.019

37. Koga, M. et al. Foxd1 is a mediator and indicator of the cell reprogramming process. Nat. Commun. (2014). doi:10.1038/ncomms4197

38. Takahashi, K. et al. Induction of pluripotency in human somatic cells via a transient state resembling primitive streak-like mesendoderm. Nat. Commun. (2014). doi:10.1038/ncomms4678

39. Gabut, M. et al. An alternative splicing switch regulates embryonic stem cell pluripotency and reprogramming. Cell (2011). doi:10.1016/j.cell.2011.08.023

40. Buganim, Y. et al. Single-cell expression analyses during cellular reprogramming reveal an early stochastic and a late hierarchic phase. Cell 150, 1209–1222 (2012).

41. Schiebinger, G. et al. Optimal-Transport Analysis of Single-Cell Gene Expression Identifies Developmental Trajectories in Reprogramming. Cell (2019). doi:10.1016/j.cell.2019.01.006

42. Guo, L. et al. Resolving Cell Fate Decisions during Somatic Cell Reprogramming by Single-Cell RNA-Seq. Mol. Cell (2019). doi:10.1016/j.molcel.2019.01.042

43. Levine, A. J. & Brivanlou, A. H. GDF3, a BMP inhibitor, regulates cell fate in stem cells and early embryos. Development 133, 209–16 (2006).

44. O’Malley, J. et al. High-resolution analysis with novel cell-surface markers identifies routes to iPS cells. Nature 499, 88–91 (2013).

45. Liu, J. et al. The oncogene c-Jun impedes somatic cell reprogramming. Nat. Cell Biol. 17, 856–867 (2015).

46. Chronis, C. et al. Cooperative Binding of Transcription Factors Orchestrates Reprogramming. Cell 168, 442–459.e20 (2017).

47. Loh, K. M. & Lim, B. A precarious balance: Pluripotency factors as lineage specifiers. Cell Stem Cell (2011). doi:10.1016/j.stem.2011.03.013

