## Supplemental Information for "Diversification of Reprogramming Trajectories Revealed by Parallel Single-cell Transcriptome and Chromatin Accessibility Sequencing"

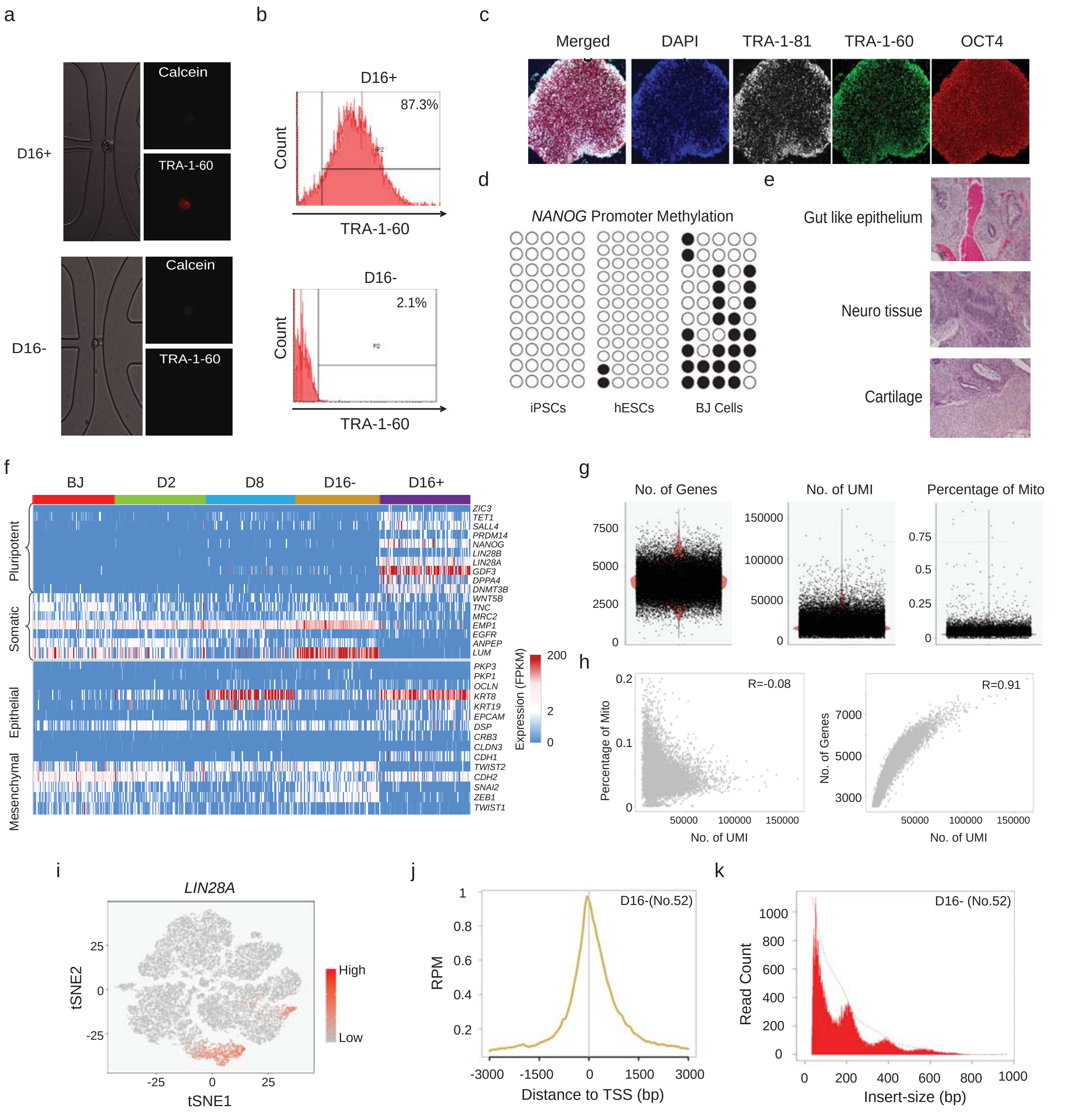


Supplementary Figure 1

**Figure S1. QC and filtering of scATAC-Seq and scRNA-Seq libraries**

(a) Representative bright field image and dye staining image including Calcein (green) and TRA-1-60 (red), in a D16+ (top) and a D16- (bottom) cell.

(b) FACS analysis of MACS sorted TRA-1-60+ (top) and TRA-1-60- (bottom) populations.

(c) Representative images of Nucleus (blue), TRA-1-81 (grey), TRA-1-60 (green) and OCT4 (red) for iPSC clones generated using polycistronic O2S, K2M reprogramming system.

(d) Bisulfite DNA methylation analysis on the promoter of NANOG in iPSCs (left), hESCs (middle) and BJ cells (right). Black circles denote methylated cytosine residue whereas the white circles denote unmethylated cytosine residues.

(e) Hematoxylin and eosin staining of teratomas derived from immunodeficient mice injected with iPSCs generated using the polycistronic O2S, K2M reprogramming system.

(f) Heatmaps demonstrating the expression levels of pluripotent and somatic genes (top), and Mesenchymal and Epithelial genes (bottom) throughout reprogramming. Each column represents a single cell, and each row indicates one gene which is listed on the right. The expression ranges from blue (low) to red (high).

(g) QC of the prepared 10x scRNA-Seq libraries demonstrating the number of genes (left), Unique molecular identifiers (UMI) (middle), and percentage of mitochondrial DNA detected (right). The 10x libraries belonging to each time-point of reprogramming were quantified using the cell ranger count module. After which, the libraries were aggregated using cellranger aggr script with disabled normalization. Next, the aggregated libraries were fed to Seurat package using the Read10X function. The QC was performed using VlnPlot function of Seurat package. The results indicate that more than 2500 genes can be detected in the majority of the 10x libraries. Additionally, most of the libraries had more than 10000 UMI. Finally, the percentage of mitochondrial DNA was found to be less than 0.25% in most of the libraries.

(h) Correlation analysis between mitochondrial DNA and UMI (left) or between number of genes and UMI (right). The analysis showed a negative correlation between UMI and mitochondrial DNA. This demonstrates that the higher number of UMI did not contribute to DNA contamination. This is further confirmed by the positive correlation between UMI and the number of genes detected. The analysis was performed using the GenePlot function of Seurat.

(i) Super-imposition of the single-cell expression levels of *LIN28A* on the tSNE plot. The Cell Range R Kit was used for this super-imposition. The tool reads the output folder of the normalized cellranger aggr script using the "load_cellranger_matrix" function. After which, unexpressed genes were filtered, UMI counts for each barcode were normalized, and the log-transformed gene-barcode matrix was calculated. Then the signature of known marker gene was visualized using the "visualize_gene_markers" function. *LIN28A* was shown to be highly expressed in cells of D8, D12 and D16+.

(j) Average enrichment profile of a D16- scATAC-Seq library around Transcription Start Sites (TSS) of the genome with a window of -3K to 3K. Y-axis denotes the average normalized read counts of the library over the indicated region in the genome (x-axis).

(k) Histograms of insert size metrics of a D16- scATAC-Seq library revealing a nucleosomal pattern which is characteristic of good ATAC-Seq library. The histogram was generated using the "CollectInsertSizeMetrics" of Picard.

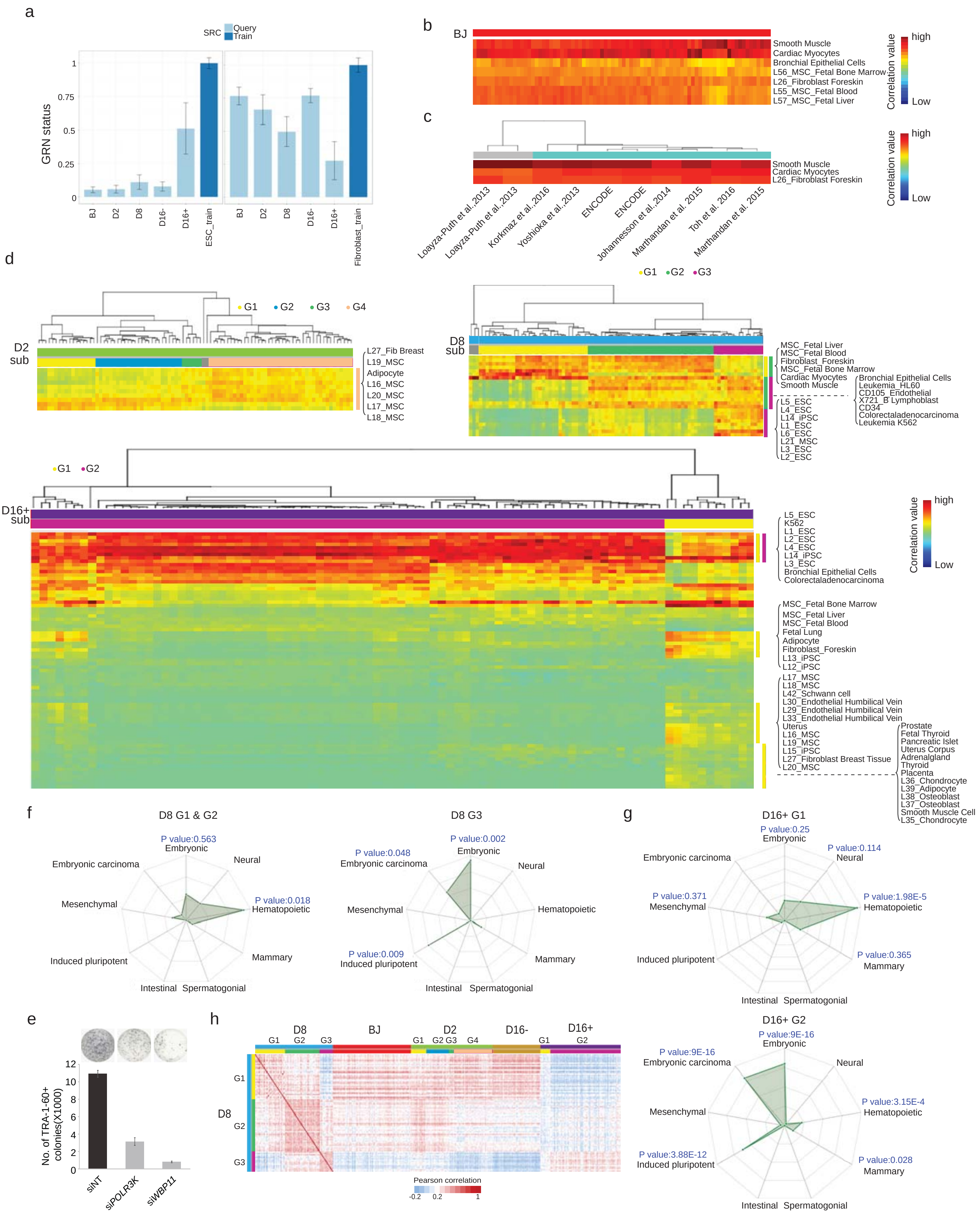


Supplementary Figure 2

**Figure S2. Classification of subgroups by RCA clustering**

(a) Bar charts exhibiting the correlation value (Y-axis) of the Fluidigm scRNA-Seq libraries of the indicated time points (X-axis) to the hESC (left) and Fibroblast (right) library from CellNet.

(b) RCA heatmap showing the highest correlated lineages of BJ scRNA-Seq libraries.

(c) RCA heatmap showing the highest correlated lineages of the published BJ bulk RNA-Seq libraries. The sources of BJ RNA-Seq libraries are indicated below.

(d) RCA heatmap showing the subgroups of D2 (top-left), D8 (top-right), and D16+ (bottom) cells clustered based on their correlation with cells of different lineage origin. Each row indicates one lineage. The correlation ranges from blue (low) to red (high). Color bar on top indicates the time points (above) and the respective subgroups (below).

(e) Bar chart demonstrating the number of TRA-1-60+ colonies (Y-axis) upon knock-down of D16+ G2 high genes, *POLR3K* and *WBP11*, at day 5 of reprogramming. Representative images are shown above. n=3. Error bar indicates SD.

(f) Stemness analysis of the highly expressed genes in D8 G1&G2 (left) and D8 G3 (right) cells. Significant adjusted P values are indicated.

(g) Stemness analysis of the highly expressed genes in D16+ G1 (top) and D16+ G2 (bottom) cells. Significant adjusted P values are presented.

(h) Heatmap of the Pearson correlation values between D8 subgroups and the cells of other time-points, based on the D8 differentially expressed genes. The correlation ranges from blue (low) to red (high). Color bar on top indicates the time points (above) and the respective subgroups (below). Side color bar indicates the subgroups of D8.

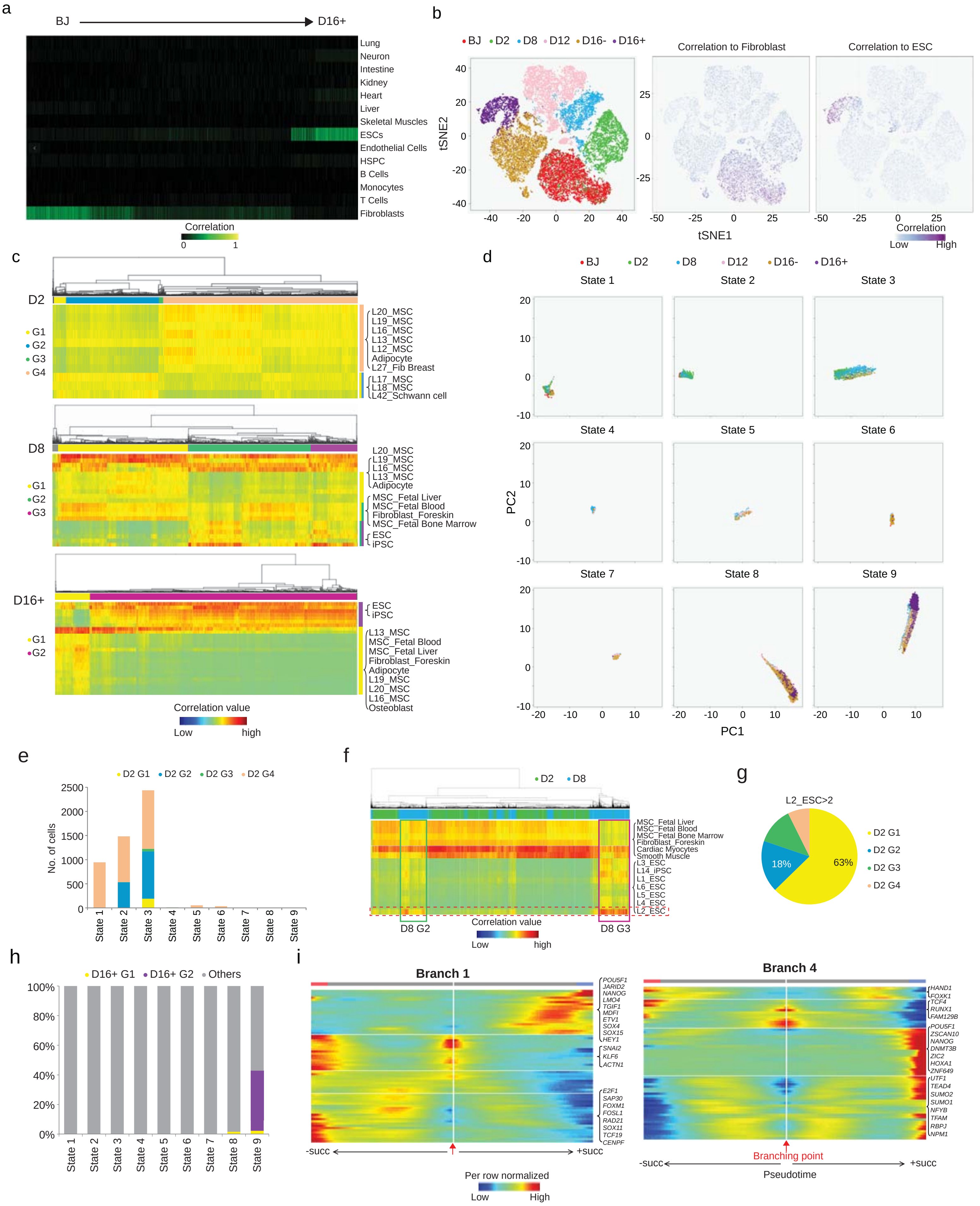


Supplementary Figure 3

**Figure S3. Reprogramming trajectories from 10x Genomics scRNA-Seq libraries**

(a) Heatmap demonstrating the correlation of the transcriptome of the 10x libraries with the global panel of CellNet. Rows represent cells of different lineages, and columns represent a 10X library. Libraries are ordered from BJ (left) to D16+ (right). Correlation ranges from black (no) to green (high).

(b) Left: tSNE clustering of the prepared 10X scRNA-Seq libraries based on the correlation to the libraries provided in CellNet. Color indicates time points. Right: Super-imposition of the correlation value of each 10X library to the Fibroblast (left) and hESC (right) on the tSNE plot. Correlation ranges from white (no) to purple (high).

(c) RCA heatmap showing the subgroups of D2 (top), D8 (middle), and D16+ (bottom) cells, clustered based on their correlation with different lineages. Each row indicates one lineage. The correlation ranges from blue (low) to red (high). Color bar on top indicates the subgroups.

(d) Trajectory states of reprogramming cells identified from the 10x scRNA-Seq libraries based on DDRTree dimension reduction. The plots indicate the time-points enriched in each state of the reprogramming trajectory.

(e) Stacked columns revealing the number of cells from the respective D2 subgroups in each reprogramming trajectory state. Color represents the subgroups of D2.

(f) RCA heatmap showing the clustering of D2 and D8 cells based on their correlation with cells of different lineage origin. Each row indicates one lineage. The correlation ranges from blue (low) to red (high). Color bar on top indicates the time points (above).

(g) Piechart revealing the distribution of D2 subgroups with the significant correlation to the RCA library “L2_ESC”.

(h) Stacked columns revealing the percentage of the indicated D16+ subgroups across the trajectory states.

(i) Heatmaps demonstrating the expression pattern of significant genes at trajectories around Branch point 1 (left) and Branch point 4 (right). Red arrow indicates the branching point. +succ denotes progress in pseudotime towards the successful trajectory with reference to the corresponding branch point. Contrastingly, -succ denotes progress towards the unsuccessful route. Expression levels ranges from dark blue (no) to dark red (high).

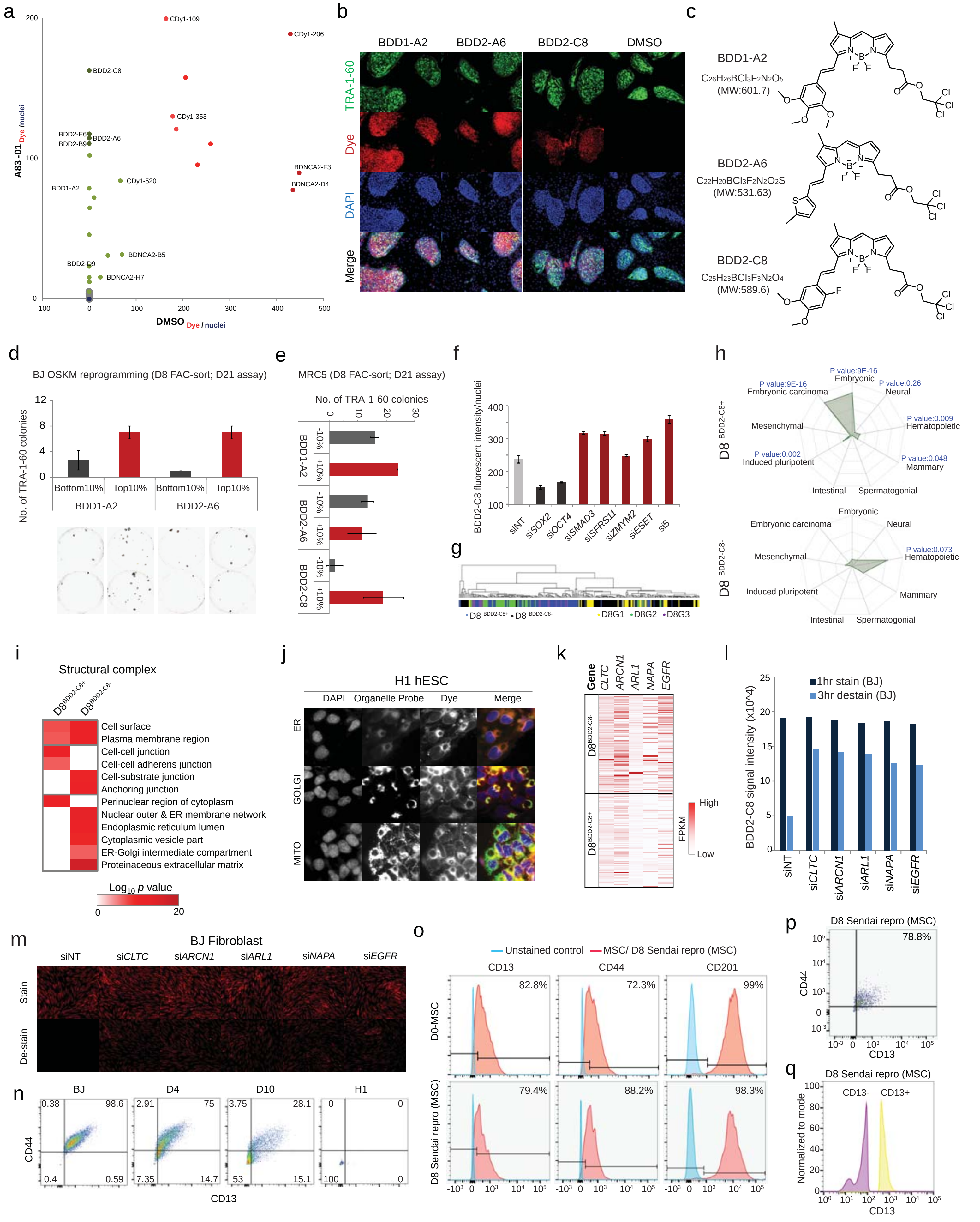


Supplementary Figure 4

**Figure S4. Identification of chemical dyes and surface markers to enrich for early reprogramming cells**

(a) Distribution plot for the chemical dyes screen. Signal was normalized to cell number (Dye/nuclei) and compared between A83-01 and DMSO control. Dyes that only stained cells in A83-01 but not DMSO control (green) were capable of distinguishing early reprogrammed cells.

(b) Representative images of staining with chemical dyes and TRA-1-60 on D21 reprogramming cells. Cells were stained with the indicated chemical dyes and subsequently fixed for TRA-1-60 immunofluorescence staining. Candidate chemical dyes co-localized with TRA-1-60 signal.

(c) Chemical structure and molecular formula of the candidate fluorescent probes.

(d) Quantification of TRA-1-60+ colonies yielded from D8 cells sorted with the respective chemical dyes (top). Top 10% and bottom 10% of the stained cells were collected and seeded for TRA-1-60 Immuno-histochemical staining at D21. Representative images are shown below. n=3; error bar indicates SD.

(e) Quantification of TRA-1-60+ colonies yielded from D8 cells sorted with the indicated chemical dyes at day 8 of reprogramming, induced from MRC5. n=2; error bar indicates SD.

(f) Barchart showing the staining intensity of BDD2-C8 in the reprogramming cells, upon depletion of the reprogramming regulators via siRNA at day 5 of reprogramming. Staining was performed at day 12. si5 refers to the combined knockdown of *SMAD3*, *SFRS11*, *ZMYM2*, *ESET* and *SAE1*. n=4; error bar indicates SD.

(g) Hierarchical dendrogram of the RCA clustering of D8 subgroups, $D8^{BDD2\text{-}C8+}$ and $D8^{BDD2\text{-}C8-}$ cells. Color represents the cell identity.

(h) Stemness analysis of the highly expressed genes in $D8^{BDD2\text{-}C8+}$ (top) and $D8^{BDD2\text{-}C8-}$ (bottom) cells. Significant adjusted P values are listed on top.

(i) Heatmap revealing the enriched structural complexes by the genes differentially expressed between $D8^{BDD2\text{-}C8+}$ and $D8^{BDD2\text{-}C8-}$ cells. Enrichment ranges from white (no) to dark red (high).

(j) Confocal images of staining with organelle-specific probes and BDD2-C8 in H1 hESCs. Green indicates organelle probes signal, red indicates dye signal, and blue indicates nuclei staining signal.

(k) Heatmap demonstrating the expression of genes involved in secretory pathway in $D8^{BDD2\text{-}C8+}$ and $D8^{BDD2\text{-}C8-}$ cells. Expression ranges from white (no) to red (high).

(l) Barchart showing the BDD2-C8 signals upon depletion of the genes involved in the secretory pathway in BJ fibroblasts. The cells were stained with BDD2-C8 at 72 hrs post siRNA mediated knockdown for FACS analysis. The knockdown of these genes did not affect the uptake of BDD2-C8 in both cell types. However, the knockdown negatively influenced the secretion and removal of BDD2-C8 from BJ fibroblasts.

(m) Representative images of the BDD2-C8 signals upon depletion of the genes involved in the secretory pathway in BJ fibroblasts. The cells were stained with BDDD2-C8 at 72 hrs post siRNA mediated knockdown. Cells were stained with BDD2-C8 for 1 hr followed by de-staining for 3 hr.

(n) Dotplots showing the co-staining signal of CD13 and CD44 in the indicated cell lines or the reprogramming cells at the indicated time points. Lines indicate the gating threshold.

(o) Overlaid histograms showing the fluorescence intensity (X-axis) of CD13 (left), CD44 (middle), and CD201 (right) in the MSC parental cells or D8 Sendai reprogramming cells induced from MSC. The numbers on top indicate the percentage of positively stained cells with the respective surface markers.

(p) Dotplots showing the co-staining signal of CD13 and CD44 in the D8 Sendai reprogramming cells induced from MSC. Lines indicate the gating threshold.

(q) Histogram showing the CD13 intensity of the sorted CD13+ and CD13- cells.

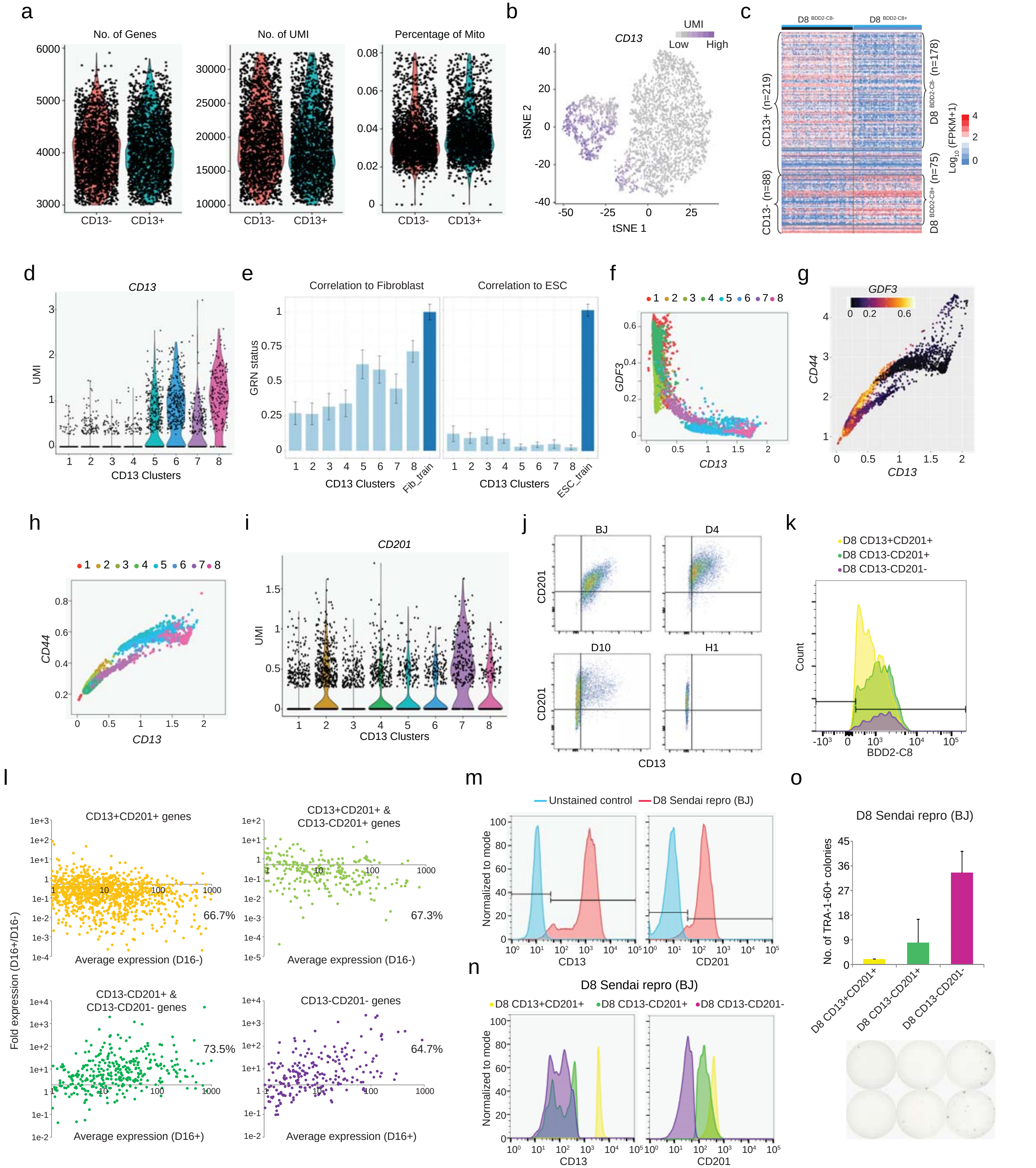


Supplementary Figure 5

**Figure S5. Analysis of CD13 10X scRNA-Seq libraries**

(a) Violin plots demonstrating the number of genes (left), Unique molecular identifiers (UMI) (middle), and percentage of mitochondrial DNA detected (right) of the D8 CD13 sorted 10x scRNA-Seq libraries.

(b) Super-imposition of *CD13* expression on the tSNE plot of the 10x D8 CD13-sorted scRNA-Seq libraries. Color represents expression level, ranging from grey (low) to purple (high).

(c) Heatmap showing the expression of the differentially expressed genes of CD13- and CD13+ in $D8^{BDD2\text{-}C8+}$ and $D8^{BDD2\text{-}C8-}$ cells. Color represents the expression level, ranging from dark blue (low) to dark red (high). The expression values are Log10 (FPKM+1).

(d) Violin plot demonstrating the expression of *CD13* across the identified CD13 sub-clusters.

(e) Bar charts exhibiting the correlation value (Y-axis) of the indicated CD13 sub-cluster (X-axis) to the Fibroblast (left) and hESC (right) library from CellNet.

(f) Super-imposition of CD13 sub-clusters on the MAGIC plot of *CD13* and *GDF3*. Color indicates the CD13 sub-clusters.

(g) MAGIC plot showing the correlative expression of *CD13* and *CD44* in the D8 CD13-sorted 10x libraries. Color represents the expression level of *GDF3*, ranging from black (no) to yellow (high).

(h) Super-imposition of CD13 sub-clusters on the magic plot of *CD13* and *CD44*. Color indicates the CD13 sub-clusters.

(i) Violin plot demonstrating the expression of *CD201* across the identified CD13 sub-clusters.

(j) Dotplots showing the co-staining signal of CD13 and CD201 in the indicated cell lines or the reprogramming cells at the indicated time points. Lines indicate the gating threshold.

(k) Overlaid histogram showing the staining intensity of BDD2-C8 in 3 populations of D8 cells with the indicated surface marker profile.

(l) Dotplots showing the comparative expression of the genes differentially expressed among the 3 sorted populations of D8, in D16+ and D16- cells. Each dot represents a gene which is highly expressed in the D8 cells with the surface markers profile indicated on top of each dotplot specifically. Y-axis represents the fold change in expression (D16+/D16-), whereas X-axis denotes the mean expression of the respective gene in either D16- (above) or D16+ (below) cells. The number besides each plot indicates the percentage of genes expressed higher (more than 2 fold) in D16- (above) or D16+ (below) cells.

(m) Overlaid histograms showing the staining intensity of CD13 (left) and CD201 (right) in the D8 Sendai reprogramming cells, induced from BJ cells.

(n) Histograms showing the fluorescence intensity (X-axis) of CD13 (left) and CD201 (right) in D8 Sendai reprogramming cells which are induced from BJ cells and possess the indicated surface markers profiles.

(o) Barcharts exhibiting the number of TRA-1-60+ colonies observed after sorting the D8 reprogramming cells (Sendai system) and categorizing the cells with the indicated surface markers profiles (x-axis) (top). The reprogramming was induced from BJ cells. Representative images are shown below. n=2; error bar indicates SD.

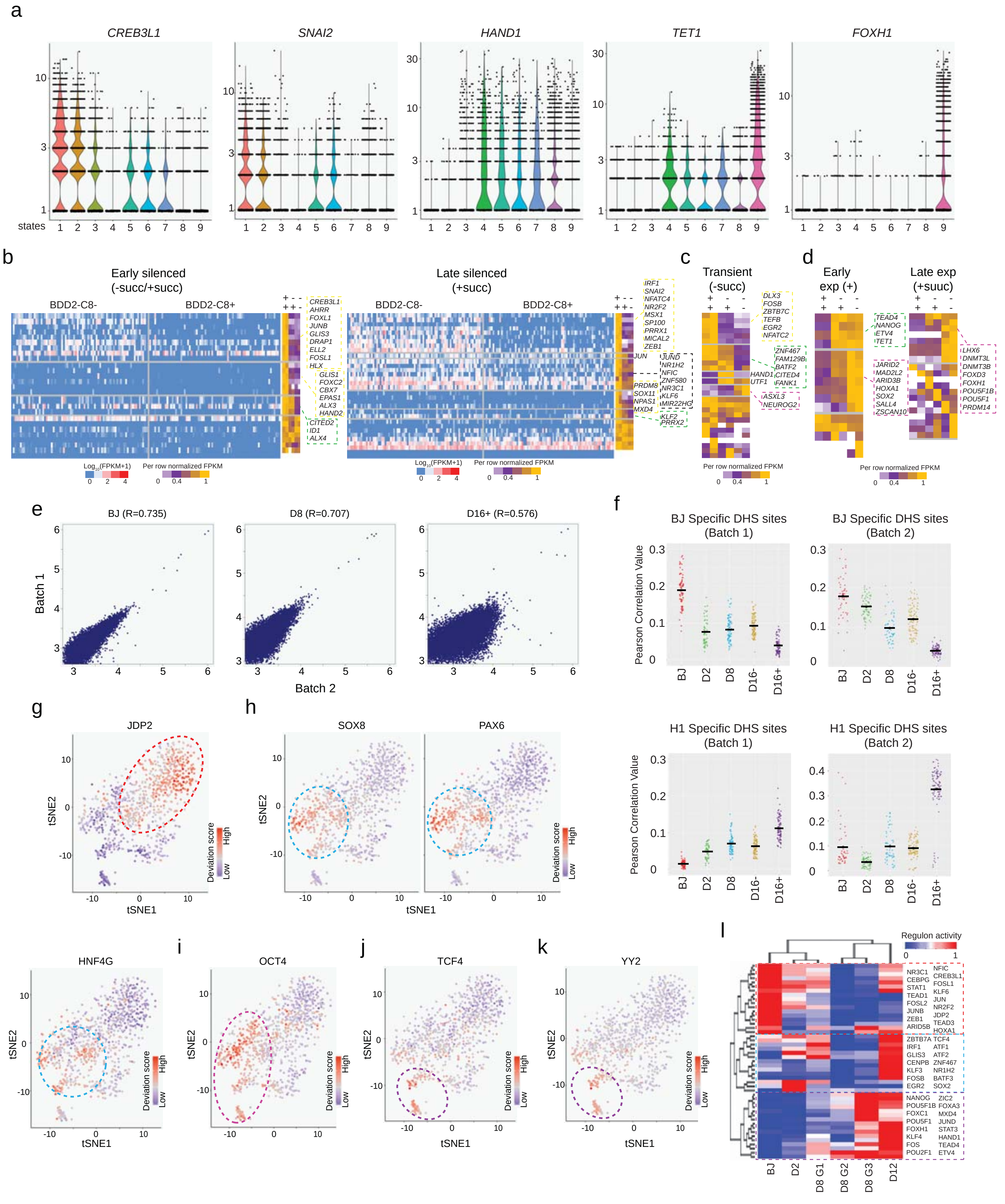


Supplementary Figure 6

**Figure S6. Classification of TFs based on the dynamics of their expression and the accessibility across reprogramming**

(a) Violin plots demonstrating the expression of the indicated TFs, belonging to the categories defined in Fig.6a, across the pseudotime states.

(b) Heatmaps showing the expression of early (left) or late (right) silenced TFs in $D8^{BDD2\text{-}C8+}$ and $D8^{BDD2\text{-}C8-}$ cells. Color represents the expression level and ranges from dark blue (low) to dark red (high expression). The expression values are Log10 (FPKM+1). Expression of these TFs in the 3 sorted populations of D8 cells is indicated beside. TFs that are highly expressed in double positive or single positive cells are labelled on the right. Color represents the per row normalized expression value of the genes and ranges from purple (low) to yellow (high).

(c) Heatmap showing the expression of the transiently expressed TFs in the 3 sorted populations of D8 cells. Differentially expressed TFs are labelled on the right. Color represents the per row normalized expression value of the genes and ranges from purple (low) to yellow (high).

(d) Heatmaps showing the expression of the early (left) and late (right) expressed TFs in the 3 sorted populations of D8 cells. TFs that are highly expressed in double negative cells are labelled on the right. Color represents the per row normalized expression value of the genes and ranges from purple (low) to yellow (high).

(e) Correlation between two replicates of scATAC-Seq libraries (batch 1 and 2) of BJ (left), D8 (middle) and D16+ (right) cells, based on the accessibility levels on predicted promoters. The axes represent the log10 of the accessibility level in each replicate.

(f) Strip charts showing the Pearson correlation values of the scATAC-Seq libraries of batch 1 and 2, against BJ (above) and H1 (below) specific DNaseI Hypersensitive Sites (DHS). Black lines represent the mean accessibility value at the indicated time-points.

(g-k) Super-imposition of the motif enrichment scores for Close motif- JDP2 (g), Transient motif-SOX8, PAX6, and HNF4G (h), and Opened motif: Early Open-OCT4 (i); Late Open-TCF4 (j) and YY2 (k) on the scATAC-Seq tSNE plot. Color indicates the accessibility level and ranges from dark blue (no enrichment) to dark red (high enrichment).

(l) Heatmap demonstrating the regulon activity of TFs calculated from the 10X scRNA-Seq libraries by SCENIC, across the indicated time points and subgroups of reprogramming. Color indicates the activity level, ranging from blue (low) to red (high).

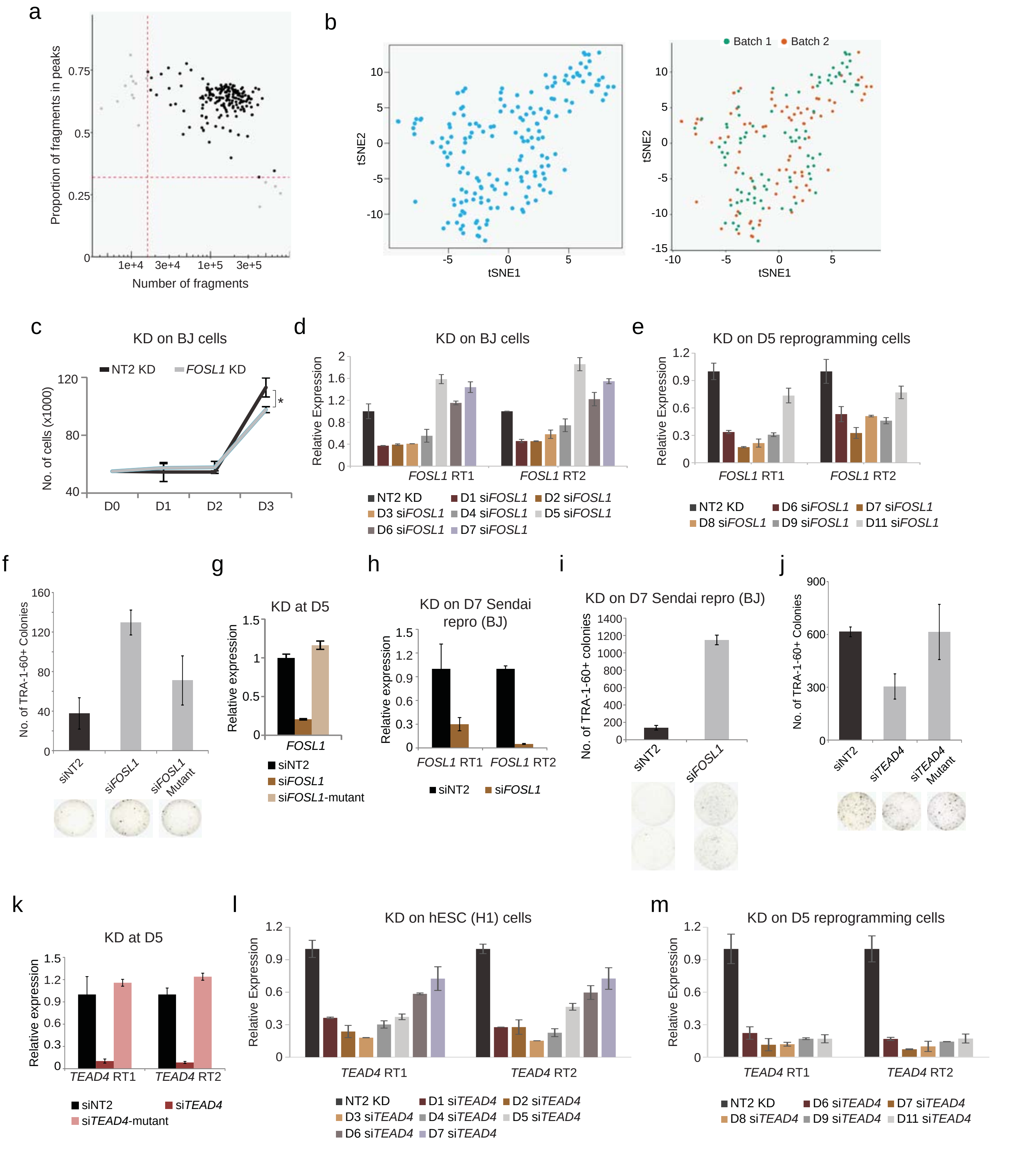


Supplementary Figure 7

**Figure S7. Functional role of FOSL1 and TEAD4 during cellular reprogramming**

(a) Dotplot revealing the proportion of fragments of each D8 scATAC-Seq library that falls in the HARs (Y-axis) plotted against the size of each library (x-axis). The red dotted lines represent the threshold for each criterion.

(b) Left: t-SNE plot of D8 scATAC-Seq libraries based on the deviation scores of JASPAR motifs. Right: Super-imposition of the batch numbers of the scATAC-Seq libraries prepared on the t-SNE plot.

(c) Line plot demonstrating the proliferation of BJ cells in which *FOSL1* was depleted (*FOSL1* KD). Non-targeting siRNA construct (NT2 KD) was used as control. X-axis denotes the time-point post siRNA transfection whereas Y-axis represents the number of cells on the respective time-point. n=3; error bar indicates SD.

(d) Bar chart exhibiting the relative expression of *FOSL1* (Y-axis) at the indicated time-points (X-axis) after depleting *FOSL1* using siRNA on BJ cells. Relative expression of *FOSL1* was calculated by normalizing to the Non-targeting siRNA constructs (siNT2) of the respective time-point. n=2. Error bar indicates SD.

(e) Bar chart exhibiting the relative expression of *FOSL1* (Y-axis) at the indicated time-points of reprogramming (X-axis) upon depleting *FOSL1* using siRNA at D5 of reprogramming. Relative expression of *FOSL1* was calculated by normalizing to the Non-targeting siRNA constructs (siNT2) of the respective time-point. n=2. Error bar indicates SD.

(f) Bar chart revealing the number of TRA-1-60+ colonies formed (Y-axis) in wells in which *FOSL1* was depleted (si*FOSL1*) at D5 of reprogramming. Mutant construct (si*FOSL1*-mutant) was used as control along with a non-targeting construct (siNT2). Representative images are shown below. n=3; error bar indicates SD.

(g) Bar chart exhibiting the relative expression of *FOSL1* (Y-axis) upon depleting *FOSL1* using wild-type (si*FOSL1*) or mutant (si*FOSL1*-mutant) siRNA constructs at D5 of reprogramming. Relative expression of *FOSL1* was calculated by normalizing to the Non-targeting siRNA constructs (siNT2). n=2. Error bar indicates SD.

(h) Bar chart exhibiting the relative expression of *FOSL1* (Y-axis) upon depleting *FOSL1* at D7 of Sendai reprogramming, induced from BJ cells. Relative expression of *FOSL1* was calculated by normalizing to the Non-targeting siRNA constructs (siNT2). n=2. Error bar indicates SD.

(i) Bar chart demonstrating the number of TRA-1-60+ colonies (y-axis) upon knock-down *FOSL1* at D7 of Sendai virus reprogramming, induced from BJ cells. Non-targeting siRNA constructs (NT2 KD) were used as controls. Representative images are shown below, n=3. Error bar indicates SD.

(j) Bar chart revealing the number of TRA-1-60+ colonies formed (Y-axis) in wells in which *TEAD4* was depleted (si*TEAD4*) at D5 of reprogramming. Mutant construct (si*TEAD4*-mutant) was used as control along with a non-targeting construct (siNT2). Representative images are shown below, n=3. Error bar indicates SD.

(k) Bar chart exhibiting the relative expression of *TEAD4* (Y-axis) at the indicated time-points (X-axis) after depleting *TEAD4* using siRNA on hESC cells. Relative expression of *TEAD4* was calculated by normalizing to the Non-targeting siRNA constructs (siNT2) of the respective time-point. n=2. Error bar indicates SD.

(l) Bar chart exhibiting the relative expression of *TEAD4* (Y-axis) at the indicated time-points of reprogramming (X-axis) when depleting *TEAD4* using siRNA at D5 of reprogramming. Relative expression of *TEAD4* was calculated by normalizing to the Non-targeting siRNA constructs (siNT2) of the respective time-point. n=2. Error bar indicates SD.

(m) Bar chart exhibiting the relative expression of *TEAD4* (Y-axis) upon depleting *TEAD4* using wild-type (si*TEAD4*) or mutant (si*TEAD4*-mutant) siRNA constructs at D5 of reprogramming. Relative expression of *TEAD4* was calculated by normalizing to the Non-targeting siRNA constructs (siNT2). n=2. Error bar indicates SD.

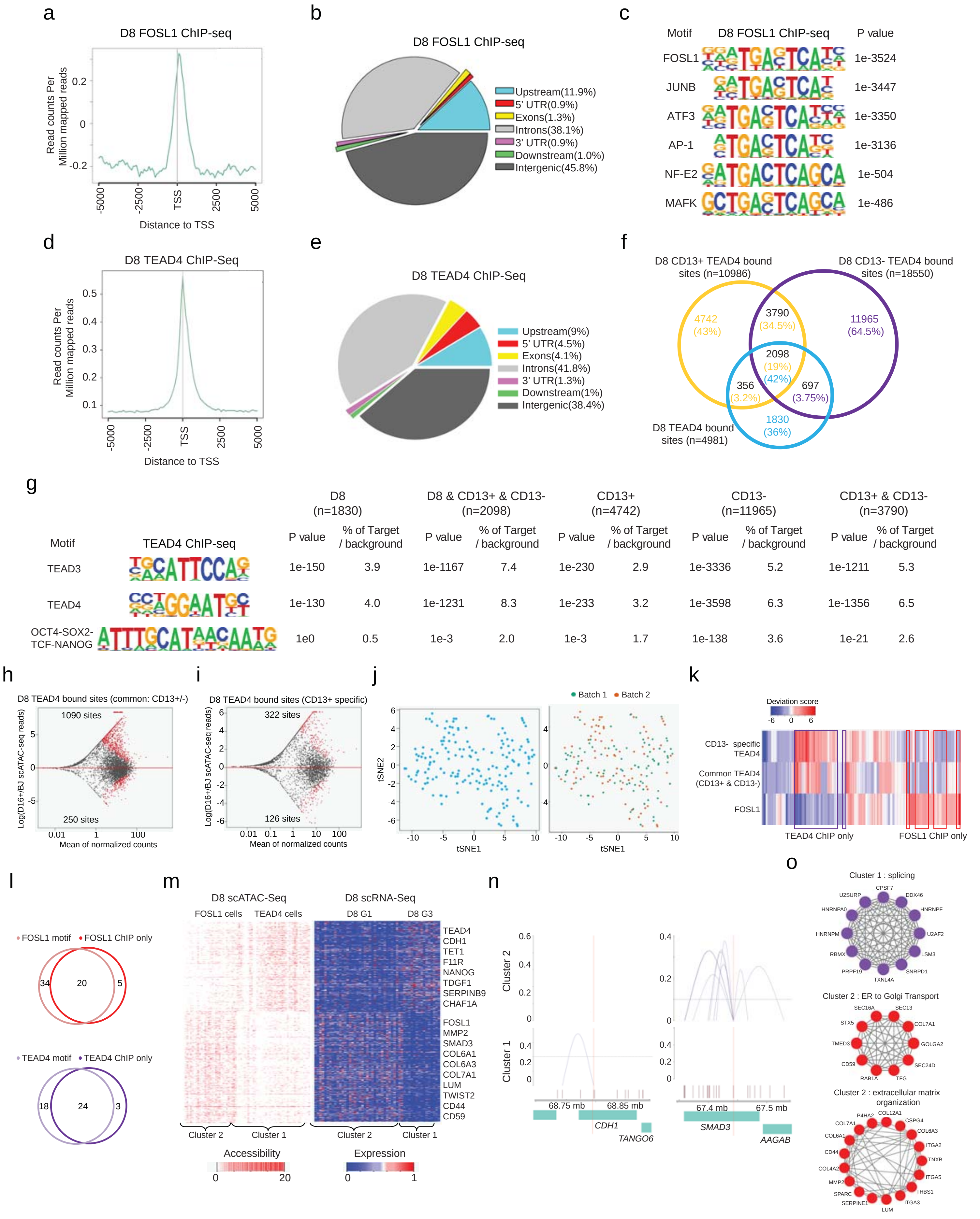


Supplementary Figure 8

**Figure S8. Genomic binding of FOSL1 and TEAD4**

(a) Average enrichment profile of D8 FOSL1 ChIP-Seq reads around transcription start sites (TSS) of the genome with a window of -5K to 5K bp.

(b) Genomic distribution of loci bound by FOSL1 in D8 reprogramming cells.

(c) Motifs significantly enriched by D8 FOSL1 ChIP-Seq sites.

(d) Average enrichment profile of D8 TEAD4 ChIP-Seq reads around all transcription start sites (TSS) of the genome with a window of -5K to 5K bp.

(e) Genomic distribution of loci bound by TEAD4 in D8 reprogramming cells.

(f) Venn diagram showing the overlap of loci detected in D8 TEAD4, D8 CD13- TEAD4, and D8 CD13+ TEAD4 ChIP-seq. The number and the percentage of bound sites falling in each category are indicated.

(g) Motifs significantly enriched by the TEAD4 bound sites detected in D8 TEAD4, CD13+ TEAD4, and CD13- TEAD4 ChIP-Seq libraries.

(h) MA plot of scATAC-Seq revealing the differentially accessible CD13+/- common TEAD4 bound sites between BJ and D16+ cells. Y-axis represents the Log Fold changes (D16+/BJ) whereas X-axis denotes the mean of normalized counts in BJ cells. Red dots denote the sites with significant accessibility changes.

(i) MA plot of scATAC-Seq revealing the differentially accessible CD13+ specific TEAD4 bound sites between BJ and D16+ cells. Y-axis represents the Log Fold changes (D16+/BJ) whereas X-axis denotes the mean of normalized counts in BJ cells. Red dots denote the sites with significant accessibility changes.

(j) Left: t-SNE plot of D8 scATAC-Seq libraries based on the deviation scores of D8 FOSL1 ChIP-seq, CD13-/+ TEAD4 ChIP-seq. Right: Super-imposition of the batch numbers of the scATAC-Seq libraries prepared on the t-SNE plot.

(k) Heatmap of D8 scATAC-Seq libraries based on the deviations scores of D8 FOSL1 specific ChIP-seq sites, CD13- specific and CD13+/- common TEAD4 specific ChIP-seq sites. Color indicates the accessibility level and ranges from dark blue (no enrichment) to dark red (high enrichment). D8 cells, showing high enrichment for FOSL1 specific ChIP-seq sites only, were labelled as “FOSL1 ChIP only” cells, whereas D8 cells, showing high enrichment for CD13- specific or CD13+/- common TEAD4 specific ChIP-seq only, were labelled as “TEAD4 ChIP only” cells.

(l) Left: Venn diagrams showing the overlap between the IDs of cells with strong enrichment of FOSL1 motif and FOSL1 ChIP only cells. Right: Venn diagram showing the overlap between the IDs of cells with strong enrichment of TEAD4 motif and TEAD4 ChIP only cells.

(m) Left: Heatmaps demonstrating the highly accessible regions (left) and highly expressed genes (right) of each cluster identified by NMF clustering. The expression heatmap (right) demonstrates the corresponding genes whose expression correlates with the accessibility of the regions in the accessibility heatmap (left). The highly accessible and expressed genes of each cluster are indicated on the right. Expression and accessibility levels range from blue (low) to red (high).

(n) Line plots indicating the Cicero co-accessibility links between the regions highlighted in red and the distal sites in the surrounding region. The height indicates the Cicero co-accessibility score between the connected peaks. The top sets of links are constructed from cells in cluster 2, while the bottom sets are built from cells in cluster 1. *CDH1* (left) is highly expressed and accessible and has more interaction with the surrounding regions in cluster 1, whereas *SMAD3* (right) are highly active and interactive in cluster 2.

(o) The interactome analysis indicating the top pathways enriched by the cluster 1 (top) and cluster 2 (middle and bottom) specific genes, which were identified from the NMF analysis.

### SUPPLEMENTAL EXPERIMENTAL PROCEDURES

#### Cell lines and reagents

Human neonatal fibroblast cell line BJ (Stemgent, Cambridge, MA) and human lung fibroblast cell line MRC-5 (ATCC ® CCL-171™) were maintained in the medium, which is composed of DME+4500mg/L medium supplemented with 10% Fetal Bovine Serum (Sigma), MEM Non-Essential Amino Acids Solution (100X), 200mM L-Glutamine (100X), 10000 U/ml Penicillin-Streptomycin (100X). Human Embryonic Stem Cells (hESCs) and induced Pluripotent Stem Cells (iPSCs) were grown in mTeSR medium (STEMCELL Technologies). The mentioned reagents were purchased from Life Technologies, unless specified.

#### Cellular reprogramming (lentivirus)

At D-1, BJ was seeded at a density of 25,000 cells/ml onto a 12-well plate. At D0, BJ cells were infected with polycistronic $O_2S$, $K_2M$ lentivirus (Addgene) in the presence of 4 mg/mL polybrene (Sigma). These cells were maintained with BJ medium until D6. At D5, cells were trypsinized, re-plated at a density of 50,000 cells/mL onto matrigel (Corning) coated 12-well plate. From D6 to D12, reprogramming cells were cultured with hESC medium, composed of 20% knockout serum replacement, MEM Non-Essential Amino Acids Solution (100X), 200 mM L-Glutamine (100X), 10,000 U/ml Penicillin-Streptomycin (100X), 8 mg/mL bFGF, 55mM β-mercaptoethanol in DMEM/F12. From day 12 onwards, medium was changed to a mixture of MEF-conditioned hESC medium and mTeSR (STEMCELL Technologies) in a 1:1 ratio.

#### Production of Viral Supernatants

293T cells were plated at a density of 1.0 x $10^7$ cells per 15cm dish. The next day, cells were transfected with 9 µg viral vector, 9 µg psPAX2 and 0.9 µg pMD2.G plasmid using 54µL TransIT LT1 (Mirus Bio) diluted with 696µL OptiMEM (Invitrogen) per plate. Supernatant was collected at

48hrs and 72hrs post-transfection, filtered through 0.45um pore size filters and concentrated using ultracentrifugation for 1.5 hrs at 23,000 RPM.

**Cellular reprogramming (Sendai virus)**

At D-2, 25k BJ or MSC cells were plated onto a 12-well plate. At D0, cells were infected with KOS, MYC, KLF4 Sendai virus (CytoTune™-iPS 2.0 Sendai Reprogramming Kit), at MOIs of 10:10:6. On the next day, medium was changed to reduce the toxicity effects of the Sendai virus. Cells were cultured in BJ and MSC medium for the first 4-5 days. At D4/5, cells were passaged and transferred onto Matrigel (Corning) coated plates. Reprogramming cells were then cultured with medium, composed of BJ/MSC medium and mTeSR medium (STEMCELL Technologies) in a 1:1 ratio. A83-01 (STEMCELL Technologies) was supplemented to the reprogramming medium to increase the reprogramming efficiency.

**Magnetic-Activated Cell Sorting (MACS)**

Enrichment of cells by MACS was performed according to the manufacturer's instructions. Briefly, cells were trypsinized to single cells and re-suspended in 1% FBS (Sigma), at a concentration of 20 million/ml. Single cell suspensions were incubated with Anti human-TRA-1-60-PE antibody, Anti human-CD13-FITC, or Anti human-CD201-PE antibody (Miltenyi Biotec) in a 1:11 dilution, at fridge for 10 min. Cells were then incubated with Anti-PE MicroBeads (Miltenyi Biotec) in a 1:5 dilution, at fridge for 15 min. Next, cells were separated into TRA-1-60 positive and TRA-1-60 negative (flow-through) cells using the magnetic sorter.

**Staining and imaging in 96-well plates (Fluorescent probes for screen)**

Cells were washed with PBS, incubated in cell culture medium with optimal concentration of fluorescent probes for 1hr, at 37$^{o}$C. It was followed by 3 washes with PBS, and destaining with culture medium for 3hrs, at 37°C. Hoechst 3342 (1:20,000, Invitrogen) was added to each well and stained for 30 min. The cells were then washed once with PBS.

Cells were imaged with an IXU ultra plate-scanning confocal microscope (Molecular Devices) at 10x magnification and 9 pictures were taken per well. Granule area, integrated fluorescent intensity and nuclei number were quantified using MetaXpress Image Acquisition and Analysis software V2.

**BDD2-C8 dye live-staining and FACS sorting**

Reprogramming cells 8d.p.i were incubated with hESC medium containing BDD2-C8 at a final concentration of 0.05 uM, at 37°C for 1hr. It was followed by 3 washes with PBS and de-staining with hESC medium for 3hrs, at 37°C. During de-staining steps, fresh hESC medium was changed every 1hr.

BDD2-C8 stained cells were trypsinized and subjected for sorting with flow cytometry (MoFlo XDP Cell Sorter, Beckman Coulter). Based on the staining intensity of BDD2-C8, top 10% and bottom 10% of D8 reprogramming cell were enriched for single cell qRT-PCR, scRNA-seq library preparation, or re-plating onto 12-well plate for TRA-1-60+ colony counting assay.

**Co-staining of cell surface markers and BDD2-C8**

Cells were stained with BDD2-C8 for 1hr, followed by 3hrs de-staining. Next, cells were trypsinized, and re-suspended with 1% FBS (Sigma) at a concentration of 1.5 million/ml. Single cell suspensions were incubated with Anti human-CD13-FITC, Anti human-CD201(EPCR)-APC-Vio770 antibody (at a dilution of 1:11) and Anti human-CD44-VioBlue (at a dilution of 1:51) (Miltenyi Biotec), at fridge for 10 min. Cells were then subjected for flow cytometry analysis.

**Organelle Probe and Fluorescent Probe Co-staining**

Cells were incubated with organelle specific probes (diluted with culture media), at 37$^{o}$C for 30min and washed with PBS 3 times. Cells were then stained with fluorescent probes at 37$^{o}$C for 1hr, followed by de-staining at 37$^{o}$C for 3hrs. The information of organelle specific probes are as follows: ER specific probe: ER-Tracker™ Red, 500nM; Golgi specific probe: BODIPY® FL C5-ceramide, 5 µM; Mitochondria specific probe:MitoTracker® Green FM, 250 nM.

**siRNA transfection in 12-well plates**

siRNA (50µL of 500nM siGENOME, Dharmacon) were prepared and frozen at -20oC before use. For reverse transfection, a master mix of 0.8µL of DharmaFECT1 (Dharmacon) transfection reagent and 150µL of OptiMEM (Invitrogen) mix was added to the tubes containing the siRNA. The DharmaFECT1 and siRNA mix was incubated for 20 min before transferring to 80,000 cells in 800µL medium to a well of 12-well plate.

**TRA-1-60 immunohistochemical staining**

Cells were fixed with 4% paraformaldehyde for 20 min at room temperature. After 3 PBS washes, cells were blocked with 7% Fetal Bovine Serum (Gibco) for 30 min at room temperature. Cells were then incubated with TRA-1-60, biotin (eBioscience, Cat. No: 13-8863-82) diluted by 1:300, at 4$^{o}$C overnight. After 3 PBS washes, cells were next stained with HRP-Streptavidin antibody (Biolegend, Cat. No: 405210) diluted by 1:500, at room temperature for 1 hr. Pierce DAB Substrate Kit (Vector laboratories, Cat. No.: SK-400) was applied for visualizing the TRA-1-60+ colonies. Plate was coated with milk and scanned by EPSON Perfection 4490 scanner. The image was analyzed by Cell Profiler.

**Immunostaining**

Cells were washed with PBS and fixed in 4% paraformaldehyde for 20 min at room temperature. After 3 PBS washes, cells were permeabilized by 0.25% Triton-X. Next, plates were washed with

PBS 3 times, and blocked in 7% Fetal Bovine Serum (Gibco) for 30 min at room temperature. Cells were then incubated with the primary antibody, at $4^{o}C$ overnight. After 3 PBS washes, cells were stained with secondary antibody for 1hr at room temperature. Hoechst 3342 (1:20,000, Invitrogen) was added to and stained for 30 min.

**Single cell qPCR**

Single cells were captured by C1 IFC for Preamp (Medium). On C1 IFC, lysis, reverse transcription with Taqman Gene Expression Assays, and pre-amplification was carried out, following *Using C1 to Capture Cells from Cell Culture and Perform Preamplification Using TaqMan Assays* manual. The pre-amplified products were harvested from C1 IFC and loaded onto 48x48 Dynamic Array IFC, along with the Taqman Gene Expression Assays for genes of interest, following *Gene Expression with the 48.48 IFC Using Fast TaqMan Assays (Biomark HD Only)* instruction. Single cell qPCR data was analyzed by Biomark HD Data Collection software.

**RNA extraction, reverse transcription and Real-time qPCR**

Total RNA was extracted using the Trizol reagent (Invitrogen). Contaminant DNA was removed by DNaseI (Ambion) treatment, and the RNA was further purified using QIAGEN RNeasy Kit. First strand cDNA was synthesized using the iScript cDNA synthesis kit (Bio-Rad) according to the manufacturer’s instructions. Quantitative real-time PCR was performed on the CXF384 Real-time System (Bio-Rad), using a Kapa SYBR Fast qPCR kit (Kapa Biosystems). The expression level of each gene was normalized to the expression level of β-actin.

**Bulk RNA-Seq**

RNA-seq libraries were prepared using the TruSeq® Stranded mRNA Library Prep kit following manual guide of the kit. Briefly, mRNA was enriched by the beads conjugated with Poly-T oligo, fragmentized, and reverse transcribed to cDNA. After ligating adaptors, cDNA was further amplified by 15 cycles of PCR. The library was sequenced on Hiseq 4000 sequencer.

**Single-cell RNA-seq (scRNA-seq) library construction**

The scRNA-seq libraries were prepared following the published protocol with minor modifications[23]. Briefly, Cells undergoing reprogramming were trypsinized into single cells at each time-point. Cells were washed 3 times in C1 DNA-seq Cell Wash Buffer (Fluidigm). Cells at a concentration of 300-350 cells/μl were combined with C1 Cell Suspension Reagent at a ratio of 3:2.5. Next, the cell mixture was loaded into the C1 Single-Cell Auto Prep IFC microfluidic chips according to the "STRT seq, 1862× (10-17 um diameter cells)" protocol. Cells were captured in the 96 capture sites in the chip and stained using a green-fluorescent calcein-AM dye (LIVE/DEAD cell viability assay, Life Technologies), and imaged by Leica CTR 6000 microscope. mRNA-Seq libraries were prepared following *Generate cDNA libraries with the C1 Single-Cell mRNA Seq HT IFC and Reagent Kit V2* manual.

**Single-cell ATAC-seq (scATAC-seq) library construction**

The scATAC-seq libraries were prepared following the published protocol with minor modifications[21]. Briefly, cells undergoing reprogramming at different time-points were trypsinized into single cells and washed 3 times in C1 DNA-seq Cell Wash Buffer (Fluidigm). Cells at a concentration of 300-350 cells/μl were combined with C1 Cell Suspension Reagent at a ratio of 3:2.5. The mixture of cells was then loaded into the C1 Single-Cell Auto Prep IFC microfluidic chips according to the "ATACseq: Cell Load and Stain (1862x)" protocol. Cells were captured in the 96 capture sites in the chip and stained using a green-fluorescent calcein-AM dye (LIVE/DEAD cell viability assay, Life Technologies), and imaged by Leica CTR 6000 microscope.

On the IFC, the lysis and transposition reaction took place at 37 $^{o}$C for 30 min, followed by inactivation of Tn5 using EDTA at 50 $^{o}$C for 30 min. Next, excess EDTA was quenched by $MgCl_2$ at room temperature. The digested accessible regions were next amplified for 8 cycles. The 96 scATAC-seq libraries were harvested from microfluidic chips and transferred to 96-well plate for incorporating cell specific indexes by additional 14 cycles of PCR. Following that, the 96 scATAC-seq libraries were pooled and purified by Qiagen PCR purification kit and quality of the libraries was assessed by Agilent 2100 bioanalyzer.

**Chromatin Immunoprecipitation (ChIP)**

ChIP was performed according to a previous study[59]. Briefly, 10 million cells were crosslinked using 1% formaldehyde for 10 min at room temperature followed by quenching by using 200 mM glycine. After that, cells were lysed using lysis buffer consisting of 10 mM Tris-Cl (pH 8), 100 mM NaCl, 10 mM EDTA, 0.25% Triton X-100 and protease inhibitor cocktail (Roche). Cells were then resuspended in 1% SDS lysis buffer composed of 50 mM HEPES-KOH (pH 7.5), 150 mM NaCl, 1% SDS, 2 mM EDTA, 1% Triton X-100, 0.1% NaDOC and protease inhibitor cocktail. The suspension was nutated for 15 mins at cold room before spinning down to collect the chromatin pellet. The pellet was then washed two times with 0.1% SDS lysis buffer containing 50 mM HEPES-KOH (pH 7.5), 150 mM NaCl, 0.1% SDS, 2 mM EDTA, 1% Triton X-100, 0.1% NaDOC, 1 mM PMSF and protease inhibitor cocktail. Chromatin was then fragmented by sonication which was conducted with cells on ice with 14 cycles, power amplitude of 35% and 30 sec pulses on with 59.9 s pulses off (Branson Sonifier 250). The chromatin solution was clarified by centrifugation at 20,000 g at 4°C for 45 mins and then pre-cleared with Dynabeads protein A (Life Technologies) for 2 hrs at 4°C. The pre-cleared chromatin sample was incubated with 50 µl of Dynabeads protein A loaded with 5 µg antibody overnight at 4°C. The beads were washed three times with 0.1% SDS lysis buffer, once with 0.1% SDS lysis buffer/0.35 M NaCl, once with 10 mM Tris-Cl (pH 8), 1 mM EDTA, 0.5% NP40, 0.25 LiCl, 0.5% NaDOC, and once with TE buffer (PH8.0). The immunoprecipitated material was eluted from the beads by heating for 45 min at 68°C in 50 mM TrisCl (pH 7.5), 10 mM EDTA, 1% SDS. Reverse crosslinking was carried out by incubating the samples with 1.5 µg/ml Pronase at 42°C for 2 hr followed by a 6 hours incubation at 67°C. DNA was then purified using MinElute PCR Purification kit (Qiagen).

Purified ChIP DNA was further processed for library preparation following Fold enrichment of each primer is calculated by normalizing to a negative control primer.

**Computational analysis:**

**Single-cell RNA-Seq libraries analysis (Fluidigm)**

Libraries were mapped to hg19 assembly using STAR aligner allowing up to 2 mismatches and removing reads mapping to more than one region. Cuffquant was used to count reads belonging to each gene in the genome with the option -u and -b included in the used script. Cuffnorm was employed to normalize the counts and calculate the FPKM among libraries belonging to the same time-point.

**Filtering single-cell RNA-Seq libraries (Fluidigm)**

Percentage of genes measures and percentage of mapping to exons were determined using the RNA-Seq QC options of SeqMonk software (https://www.bioinformatics.babraham.ac.uk/projects/seqmonk/). Libraries with mapping to exon rate below 75% were filtered out. Moreover, libraries with below 20% of measured genes were also filtered out.

**Distribution of reads over transcripts (Fluidigm)**

The determination of the distribution of reads over the transcripts of house-keeping genes was performed using the geneBody_coverage.py module of RseQC python package.

**Differential expression analysis (Fluidigm scRNA-Seq)**

The differentially expressed genes were calculated among subgroups of each time point. If the gene (1) has average FPKM value more than 5, (2) was expressed in more than 50% of the cells, (3) was expressed as high as more than 3 fold in average expression in the subgroup, it is considered to be highly expressed in the respective subgroup.

**Gene ontology analysis**

DAVID (https://david.ncifcrf.gov/) was used for performing gene ontology analysis. The GO terms were extracted from GOTERM_BP_DIRECT under the Functional Annotation tab.

**Boxplot**

Boxplots were generated by "ggplot2" package in R, based on the $Log_{10}$ (FPKM+1) value of specified genes in all the subgroups.

**Pearson correlation**

Pearson correlation between subgroups of D8/D16+ and cells from all the other time points was performed using EXCEL, based on the differentially expressed genes of D8/D16+.

**Differential expression analysis (10X Genomics scRNA-Seq)**

The differentially expressed genes were calculated among subgroups of each time point. If the gene (1) has average UMI value more than 0.2, (2) was expressed in more than 15% of the cells, (3) was expressed as high as more than 3 fold in average expression in the subgroup, it is considered to be highly expressed in the respective subgroup. Due to the depth of 10X Genomics scRNA-Seq libraries, the threshold is different from Fluidigm scRNA-Seq libraires.

**STEMNESS score analysis**

Stemness score was obtained by inputting the differentially expressed genes of each subgroup in Stem Checker (http://stemchecker.sysbiolab.eu/).

**D8 Subgroups Classifier creation (Fluidigm)**

Machine learning was performed using "MLSeq"[60]. Briefly, the mapped bam files were sorted by "read name" using samtools sort with option -n. The sorted files were then used as an input for htseq-count feature of HTSeq[61] to obtain raw gene counts. The option strand was set to reverse. The output table of HTSeq was used as an input for MLSeq. First we trained the machine using raw matrix of D8 cells in which the subgroups were pre-determined using RCA. The input was converted to DESeq matrix using "DESeqDataSetFromMatrix" option from DESeq2[62]. RandomForest was the algorithm of choice for training. The training was performed using the following script "classify(data = data.trainS4, method = "rf", preProcessing = "deseq-vst", ref = "G1", tuneLength = 10, control = trainControl(method = "repeatedcv", number = 5, repeats = 10, classProbs = TRUE))".

After the training was completed, we used the entire raw matrix as input. The cells in the matrix were annotated according to their time-point. Next, the matrix was converted into DESeq matrix as mentioned above. Predictions were performed using "predict" finction of MLSeq. D8 cells were used to test the accuracy of the classifier as the subgroups of D8 cells were already known.

**QC of 10x Genomics libraries analysis**

Fastqs were generated from the 10x raw data using the mkfastq module of cellranger. The 10x libraries belonging to each time-point of reprogramming were quantified using the cell ranger count module. Then, the libraries were aggregated using cellranger aggr script with disabled normalization by setting the option --normalize=NONE. Next, the aggregated libraries were fed to Seurat package using the Read10X function. The QC was performed using VlnPlot function of Seurat package. Mitochondrial DNA percentage in each library was measured by grepping genes starting with MT and summing the values using colSums. The correlation between number of genes and UMI or between percentage of Mitochondrial DNA and UMI was performed using the GenePlot function of Seurat. Cells with detected genes below 2500 or above 6500 were filtered out.

**tSNE Clustering of 10X libraries**

After the quantification of the 10x libraries had been performed using cellranger count, the libraries belonging to different time-points were aggregated using the cellranger aggr function with mapping-based normalization. This ensured that all the libraries had equal number of confidently mapped reads. The script provided tSNE plot after its completion. For super-imposing the expression of genes on these clusters, the Cell Range R Kit was used. The tool reads the output folder of the normalized cellranger aggr script using the "load_cellranger_matrix" function. After which, unexpressed genes were filtered, UMI counts for each barcode were normalized, and the log-transformed gene-barcode matrix was calculated. Then the signature of known marker genes was visualized using the "visualize_gene_markers" function.

**CellNet**

For Fliudigm scRNA-Seq libraries, the mapped bam files were sorted by "read name" using samtools sort with option -n. The sorted files were then used as an input for htseq-count feature of HTSeq[61] to obtain raw gene counts. The option strand was set to reverse. The output table of HTSeq was used as an input for CellNet[24].

For 10X Genomics scRNA-Seq libraries, the "mat2csv" function of cellranger was used to generate a raw gene count matrix from the "cellranger aggr" ouput folder (raw) with Gene Symbols. This matrix was used as an input for CellNet.

The analysis was performed by converting the matrices to rda files and uploaded to CellNet using the utils_loadObject option. The data was then transformed by using weighted_down (1e3) and trans_prop options of cell net. The trained data and Salmon index tables used for human were downloaded from https://github.com/pcahan1/CellNet. The training of our scRNA-Seq libraries was performed using the cn_apply function as instructed in CellNet manual. The tSNE was generated by finding variable genes using findVarGenes and then calculating PCA and tsne using prcomp and to_tsne as instructed by CellNet Manual. Bar plots were generated using cn_barplot_grnSing and the heatmap was generated using cm_HmCalls.

**Pseudotemporal analysis**

The output folder of cellranger aggr (without normalization) was fed into Monocle 2 using the newCellDataSet function. The "negbinomal.size" expression family was chosen. After which the size factors and dispersions of the libraries were estimated. Genes that were expressed in minimum of 10 cells were considered for subsequent analyses. Differential gene was performed using "differentialGeneTest" while using the size factors for correction of the estimations. Genes which are differentialy expressed with qval < 0.01 were considered for the tree constructions. Dimension reduction was performed using DDRTree methods and the trajectory was plotted using the plot_cell_trajectory. For super-imposition of genes expression on the measured cell trajectories, the marker option of plot_cell_trajectory option was used and use_color_gradient was set as TRUE.

**Branching analysis**

All the branches in single cell trajectories were analyzed using BEAM. The analysis enabled us to identify the branch-dependent gene expression profile at the single-cell level. Genes which were shown to be exclusively expressed in branches associated with unsuccessful reprogramming for each branch point (q-value = 0) were selected for the analysis. The genes were uploaded to DAVID and the "Biological Processes Direct" tab was selected. For the generation of branching heatmaps, genes of interest were made in to a dataframe and this data frame was used in the "plot_genes_branched_heatmap" script of monocle.

**Determination of Regulatory TFs**

Differentially expressed genes across the States were determined using "differentialGeneTest" function. The "fullModelFormulaStr" was set to States. The genes which were significant were then annotated (Molecular Function) using Metascape[63] (http://metascape.org/gp/index.html). Significant TFs were then visually inspected one-by-one to determine the significant regulators across the pseudotime states.

**Seurat Analysis (10x Reprogramming Cells)**

After filtering low quality cells as indicated above, the states to which each reprogramming cell belonged to in the pseudotime trajectory were added using the addMetaData function. The TFs heatmap was generated using the "DoHeatmap" function and grouping the cells by States.

**Seurat Analysis (10x D8 CD13 sorted cells)**

CD13+ and CD13- libraries were aggregated using cellranger aggr script with disabled normalization by setting the option --normalize=NONE. Next, the aggregated libraries were fed to Seurat package using the Read10X function. The QC was performed using VlnPlot function of Seurat package. Mitochondrial DNA percentage in each library was measured by grepping genes starting with MT and summing the values using colSums. The percentage of Mitochondrial DNA was then added as a

MetaData. Cells that passed QC had genes detected between 3000-6000, UMI count between 10000-32500 and mitochondrial contamination below 0.08. Variable genes were detected using the following parameters in "FindVariableGenes" function: mean.function = ExpMean, dispersion.function = LogVMR, x.low.cutoff = 0.0125, x.high.cutoff = 3, y.cutoff = 0.5).

Libraries were then normalized using LogNormalize method with 10000 scale. After that, the libraries were scale to UMI and percentage of mitochondrial DNA. CD13- cells that demonstrated expression of CD13 were considered as sorting error and filtered out. PCA was determined using variable genes, and tSNE was performed using dims 1:9. Clusters were determined using FindClusters with 0.6 resolution.

**Pseudotime Analysis (Reprogramming Cells with D8 CD13 sorted cells)**

BJ, D2, D12, D16-, D16+, D8 CD13- and D8 CD13+ 10xs libraries were aggregated using cellranger "aggr" without normalization. Next, the aggregated libraries were fed to Seurat package using the Read10X function. Cells with genes detected below 1875 or above 7000 were filtered out. Additionally cells with mitochondrial DNA above 0.1 were filtered out. The Seurat object was then fed into Monocle v3 using "importCDs". Then "estimateSizeFactors" and "estimateDispersions" were performed. This was followed by "preprocessCDS" with 20 dimensions. Then dimensions were reduced using UMAP [64]. Cells were then portioned using "PartitionCells" with default parameters. Finally "learnGraph" was used "RGE_method" set to DDRTree and trajectory was plotted using "plot_cell_trajectory".

**Mathematical Imputation of 10x Libraries**

Raw expression matrix was generated using the mat2csv function of cellranger. The matrix was then transposed using R. This matrix was then used as input for MAGIC[36]. We filtered genes that had below 10 UMI and filtered cells with total UMI < 7000. The matrix was then normalized using "library.size.normalize" function of MAGIC and then transformed using "sqrt" function. Imputation was then performed using "magic" function with default parameters.

**Mapping of scATAC-Seq libraries**

Libraries were mapped to hg19 genome assembly using STAR aligner. Reads that contain more than 2 mismatches and/or mapped to more than one position in the genome were filtered out. The option --alignIntronMax was set to 1 to disable the splice awareness of the aligner. The option -alignEndsType was set to EndToEnd to make STAR compatible with DNA reads.

**Determination of Highly Accessible Regions (HARs)**

Mapped files of libraries belonging to the same time-point were merged using samtools. After that the MarkDuplicates module of PICARD tools (http://broadinstitute.github.io/picard) was used to remove duplicates. Then MACS2 was used to call peaks. The --nomodel --nolambda --keep-dup all --call-summits options were used. ChrM and all ambiguous chromosomes were excluded from this analysis. The top 50000 peaks in the peaks summit file were extended to -/+ 250 bp. These were then considered to be the HARs in each respective time-point.

**Filtering single-cell ATAC-Seq libraries**

Duplicate were removed from each single-cell ATAC-Seq as mentioned above. This was followed by the addition of group information using the AddOrReplaceReadGroups module of PICARD. A sequence dictionary for hg19 genomes lacking ChrM and other ambiguous chromosomes was generated using CreateSequenceDictionary module of PICARD. Reads in each library were sorted lexicographically by using the ReorderSam module of PICARD tools. The option ALLOW_INCOMPLETE_DICT_CONCORDANCE was set to TRUE which helps in removing ChrM and ambiguous mapped reads from the libraries. The generated bam files were then indexed using samtools. The coverage of each single-cell ATAC-Seq over the HARs of their corresponding time-point was measured using the DepthOfCoverage module of GATK tools v3.5. The option –countType was set to COUNT_FRAGMENTS to ensure reliable estimates. Libraries should have coverage over minimum 15% of the HARs coordinates to be considered for further analysis. The coverage values are recorded in the total_cvg columns present in the interval_summary output of GATK. Additionally, library size of each library was determined using the MarkDuplicates of PICARD tools. Any library having a size less than 10000 was discarded.

**Average enrichment profile generation**

The processed scATAC-seq libraries, in which ChrM and ambiguously mapped reads were removed, were used to generate the average enrichment profile plots. The software ngs.plot was used. The options -R was set to tss and -L was set to 3000 to measure the average enrichment of reads on +/-3K around the TSS regions.

**Nucleosomal pattern determination**

The CollectInsertSizeMetrics module of PICARD tools was employed to generate the insert-size histograms. The libraries were free from reads mapping to ChrM or other ambiguous chromosomes prior to running the scripts.

**UCSC genome browser screenshots**

The bedGraph file for each time-point was generated by first merging the bam files of the corresponding time-point using samtools. Secondly, HOMER's makeUCSCfile script was used with the default parameters for the scATAC-Seq and ChIP-Seq libraries. In case of scRNA-Seq libraries, the options -style and -fragLength were set to rnaseq and given respectively.

**Motif analysis**

Motifs enriched in regions of interest were determined using the findMotifsGenome.pl script of HOMER. The option size was set to "given" in the script.

**Prediction of promoters**

First, the HARs were annotated using annotatePeaks.pl script. Then, regions that were 3K upstream to 200 bp downstream from the nearest gene were considered to be putative promoters. We ensured that these regions do not fall in the genebody of any gene. The coverage of the single-cell single ATAC-seq libraries over these putative promoters was measured using GATK tools as mentioned above. The values were normalized by dividing each value present in the total_cvg column by the respective library size and multiplying that by 10000. These normalized values were then uploaded to R and PCA plots were generated using "FactoMineR".

**Differential accessibility analysis**

The differential accessibility analysis was performed using DESeq2. The unnormalized coverage values over regions of interest were fed to R. The time-point of the libraries were considered as conditions. The size factors were estimated using the "poscounts" method. This was followed by normalization and differential accessibility analysis using the "DESeq" function. The reported regions underwent Independent Hypothesis Weighting to ensure the removal of false positive hits by using the filterFun=ihw in the "results" option of DESeq2. The MA plot was generated using the plotMA function of DESeq2.

**ChromVAR analysis of scATAC-Seq libraries**

De-duplicated mapped scATAC-Seq files and the narrowPeaks HARs were uploaded to chromVAR software and the coverage of the libraries over the HARs was measured using getCounts function of chromVAR. Libraries were filtered to remove dead or double cells libraries using filterSamples option. The HARs were annotated with JASPAR motifs using matchMotifs option. Variability of

JASPAR motifs across all the cells was quantified using the computeDeviations function. This was followed by computing the variability using computeVariability option. Then, the scATAC-Seq libraries were correlated based on the computed deviations using the "getSampleCorrelation" module.

Clustering of scATAC-Seq libraries based on deviation scores which were quantified was measured using "deviationsTSNE" of chromVAR with thresholds and perplexities that ensured that the results were not due to batch effect. Super-imposition of motif enrichment score for FRA1, OCT4 and TEAD4 on the scATAC-Seq clusters. These genes were considered in the annotation field in the "plotDeviationsTSNE" script of chromVAR.

**ChIP-seq analysis**

Libraries were mapped to hg19 genome using STAR as performed for the scATAC-Seq libraries (mentioned above). Subsequently, tag directories were created for each mapped file using makeTagDirectory script of Homer. This was followed with peak calling using GEM[65]. The --k_min 6 --k_max 13 --t 10 --outHOMER –outBED were the additional options passed to GEM. The default read distribution file of GEM was used.

**Genomic distribution plots**

The identified binding sites of FRA1 were uploaded to PAVIS (https://manticore.niehs.nih.gov/pavis2/). The hg19 known Genes Assembly was used for annotating the peaks. For other parameters, the default values were used.

**Sequencing data**

Raw data is deposited in GEO with the accession number: GSE100345.

**REFERENCE**


59. Yang, B. X. *et al.* Systematic Identification of Factors for Provirus Silencing in Embryonic Stem Cells. *Cell* **163**, 230–245 (2015).

60. Zararsiz, G. *et al.* A comprehensive simulation study on classification of RNA-Seq data. *PLoS One* (2017). doi:10.1371/journal.pone.0182507

61. Anders, S., Pyl, P. T. & Huber, W. HTSeq-A Python framework to work with high-throughput sequencing data. *Bioinformatics* **31**, 166–169 (2015).

62. Love, M. I., Huber, W. & Anders, S. Moderated estimation of fold change and dispersion for RNA-seq data with DESeq2. *Genome Biol.* **15**, (2014).

63. Zhou, Y. *et al.* Metascape provides a biologist-oriented resource for the analysis of systems-level datasets. *Nat. Commun.* (2019). doi:10.1038/s41467-019-09234-6

64. McInnes, L., Healy, J., Saul, N. & Großberger, L. UMAP: Uniform Manifold Approximation and Projection. *J. Open Source Softw.* (2018). doi:10.21105/joss.00861

65. Guo, Y., Mahony, S. & Gifford, D. K. High Resolution Genome Wide Binding Event Finding and Motif Discovery Reveals Transcription Factor Spatial Binding Constraints. *PLoS Comput. Biol.* (2012). doi:10.1371/journal.pcbi.1002638
